# Hemodynamics Affects Factor XI/XII Anticoagulation Efficacy in Patient-Specific Left Atrial Models

**DOI:** 10.1101/2024.08.27.609969

**Authors:** M. Guerrero-Hurtado, M. Garcia-Villalba, A. Gonzalo, E. Durán, P. Martinez-Legazpi, A. M. Kahn, M. Y. Chen, E. McVeigh, J. Bermejo, J. C. del Álamo, O. Flores

**Affiliations:** Department of Aerospace Engineering, Universidad Carlos III de Madrid, Leganés, Spain; Institute of Fluid Mechanics and Heat Transfer, TU Wien, 1060 Vienna, Austria; Department of Mechanical Engineering, University of Washington, Seattle, WA, USA; Department of Mechanical, Thermal and Fluids Engineering, Universidad de Málaga, Málaga, Spain; Dept. of Mathematical Physics and Fluids, Universidad Nacional de Educación a Distancia, Spain; CIBERCV, Madrid, Spain; Division of Cardiovascular Medicine, University of California San Diego, La Jolla, CA, USA; National Heart, Lung, and Blood Institute, National Institutes of Health, Bethesda, MD, USA; Department of Bioengineering, University of California San Diego, La Jolla, CA, USA; Department of Radiology, University of California San Diego, La Jolla, CA, USA; Hospital General Universitario Gregorio Marañón, Madrid, Spain; Instituto de Investigación Sanitaria Gregorio Marañón, Madrid, Spain; Facultad de Medicina, Universidad Complutense de Madrid, Madrid, Spain; Center for Cardiovascular Biology, University of Washington, Seattle, WA, USA; Division of Cardiology, University of Washington, Seattle, WA, USA

**Keywords:** digital twins, coagulation cascade, factor XI, factor XII, computational fluid dynamics

## Abstract

Atrial fibrillation (AF) disrupts the circulation of blood through the left atrium (LA), and may result in relative stasis in the left atrial appendage (LAA), increasing thromboembolic risk. Anticoagulant agents can lower this risk, but currently used agents target the common pathway central to the coagulation cascade, increasing bleeding risk. Anticoagulants such as factor XI/XII inhibitors target the initial phase of the intrinsic pathway, with a significantly lower associated bleeding risk. However, these agents’ efficacy in preventing thrombosis in patient-specific flow conditions is not fully understood. We hypothesized that patient-specific flow patterns in the LA and LAA not only influence the risk of thrombosis but also the effectiveness of anticoagulation agents. We simulated blood flow and the intrinsic coagulation pathway in patient-specific LA anatomies with and without factor XI/XII inhibition to test this hypothesis. We considered thirteen patients in sinus rhythm and AF, several of whom had an LAA clot or a history of transient ischemic attacks. We used computational fluid dynamics based on 4D CT imaging and a detailed 32-species coagulation system to run 247 simulations for 13 patients, systematically sweeping over a wide range of factor XI/XII inhibition levels. Implementing a novel multi-fidelity coagulation modeling approach accelerated computations by two orders of magnitude, enabling the large number of simulations performed. Our simulations provide spatiotemporally resolved maps of thrombin concentration throughout the LA, showing it peaks inside the LAA. Coagulation metrics based on peak LAA thrombin dynamics suggested patients could be classified as *non-coagulating, moderately* and *severely coagulating* cases. *Severely coagulating* cases had significantly slower flow and higher residence time than *moderately coagulating* patients inside the LAA, requiring stronger factor XI/XII inhibition to blunt thrombin growth. The methodology outlined in this study has the potential to enable personalized assessments of coagulation risk and tailor anticoagulation therapy based on medical imaging.

## 1 Introduction

Atrial fibrillation (AF) is a common arrhythmia affecting between 20% and 33% of individuals older than 45 during their lifetime [Linz et al., 2024]. In AF, the cyclic contraction of the atria is replaced by a rapid yet erratic and weak trembling motion that disturbs flow inside these chambers. AF-associated flow is particularly aberrant inside the left atrial appendage (LAA), a small, narrow sac protruding the left atrium (LA), where thrombosis is most likely in AF patients.

Direct oral anticoagulants (DOACs) are an effective therapy for patients with AF who need long-term anticoagulation to reduce the risk of ischemic stroke and other embolic events associated with LAA thrombosis. DOACs have notable advantages over warfarin, including reduced monitoring requirements and fewer drug and food interactions [Grzésk et al., 2021]. Except for dabigatran, which is a direct thrombin inhibitor, currently used DOACs target thrombin amplification by reducing the concentration of activated factor X [Heestermans et al., 2022]. This factor has a critical function in coagulation as it connects the extrinsic pathway, externally triggered by vascular injury, with the self-initiated, intrinsic pathway [Davie et al., 1991]. Because of its central role in the coagulation cascade, factor X inhibition lowers thromboembolic risk but also increases the likelihood of internal hemorrhage, and DOAC prescription must balance these two probabilities. Existing tools to predict thromboembolic stroke risk in AF (e.g., the CHA_2_DS_2_-VASc score) have moderate accuracy [Siddiqi et al., 2022]. Moreover, the response to DOACs varies with factors like age, genetics, and metabolism [Gong and Kim, 2013]. For patients at high risk of bleeding prescribing reduced DOAC doses is sometimes done, but there are limited data evaluating the efficacy of this approach [Halperin and Rashed, 2021].

Given the complexity of personalizing DOAC prescription in AF patients, recent efforts have focused on inhibiting coagulation factors other than factor X [Campello et al., 2022]. Prior research has identified intrinsic pathway activation, mediated by factor XI/XII among other coagulation factors, as a therapeutic target with potential to reduce bleeding risk [Wilbs et al., 2020]. *In vitro* and *in vivo* experiments have tested factor XI/XII antagonists, showing that 70-90% inhibition can delay clotting time via the intrinsic pathway (i.e., measured with the activated Partial Thromboplastin Time, aPTT) without affecting clotting time via the extrinsic pathway (i.e., measured with the Prothrombin Time, PT) [Wilbs et al., 2020, Liu et al., 2011, Naito et al., 2021]. Factor XI antagonists have been proven significantly safer than currently used DOACs [Piccini et al., 2022] but their relative efficacy is still uncertain [Presume et al., 2024]. Overall, the increasing availability of drugs targeting different coagulation factors and their dose dependence create a need for improved models to understand their mechanism of action [Bikdeli et al., 2022].

Quantifying LA flow patterns and understanding how they affect the efficacy of anticoagulation agents are significant barriers towards personalizing these agents’ prescription. The idea that flow patterns affect thrombosis is universally accepted as a pillar of the Virchov’s triad [Ding et al., 2020]. Hence, it seems plausible that the effects of inhibiting coagulation factors will also be sensitive to patient-specific flow patterns. Coagulation tests measure the kinetics of relevant coagulation cascade species (e.g., thrombin, prothrombin, activated partial thromboplastin) using laboratory test kits that do not reproduce flow conditions, let alone patient-specific ones. Therefore, the accuracy of these tests’ results may thus vary from patient to patient.

*In vitro* and *in vivo* experiments of thrombosis in microvascular settings can provide highly detailed information about the underlying molecular mechanisms and the effects of flow [Welsh et al., 2014]. Accessing similar information in large-animal models is significantly more challenging, motivating the development of computational models. However, compared to other areas of patient-specific computational modeling, progress in the simulation of the coagulation cascade has been limited due to the unmanageable cost of the simulations. Coagulation under flow is mathematically modelled by a system of coupled advection-reaction-diffusion partial differential equations (PDEs), one for each reacting species, whose numerical solution is traditionally implemented within a computational fluid dynamics (CFD) solver jointly with the solution to the Navier-Stokes equations governing blood flow [Cito et al., 2013]. The large number of species involved, the fine 3D meshes required to resolve spatial gradients of species concentration, and the disparate timescales governing coagulation and blood flow [Wootton and Ku, 1999], have frustrated researchers for decades. Most previous efforts have focused on simplified geometries representative of implanted medical devices [Méndez Rojano et al., 2018] or cardiovascular anatomies [Biasetti et al., 2012, Seo et al., 2016, Qureshi et al., 2021]. More recent studies are beginning to consider patient-specific anatomies [Grande Gutiérrez et al., 2021, Qureshi et al., 2023] in limited number of configurations, none of which include anticoagulation agents. As far as we know, there are no comprehensive simulation studies on intracardiac coagulation considering multiple patients, and much less each patient’s reaction to varying anticoagulant dosages

We report results from an extensive set of simulations of LA coagulation versus factor XI/XII inhibition in 3D patient-specific models. We used CFD analysis and multi-fidelity (MuFi) coagulation modeling [Guerrero-Hurtado et al., 2023], a novel approach that decouples the blood flow and coagulation solvers, accelerating our simulations by two orders of magnitude. We considered 13 patient-specific LA models including cases in sinus rhythm and AF, patients with and without LAA clots or cerebrovascular accidents, and patients following different anticoagulation regimes. Overall, we performed 247 simulations considering 32 coagulation species and 19 levels of factor XI/XII inhibition per patient. Thrombin levels were highest in the LAA of patients with poor blood washout, and the effectiveness of new anticoagulants targeting the intrinsic coagulation pathway was also worst for these patients. The new computational tools presented in this manuscript could open new venues to improve patient selection and tailor anticoagulant prescription in AF.

The manuscript is organized as follows. The methodology is presented in section 2, including a brief description of the MuFi model, the coagulation cascade kinetics used in the models, and the patient-specific CFD simulations. Section 3 includes the verification of the MuFi models for patient-specific LA flow, the description of the thrombin concentrations without inhibition, and the effects of factors XI/XII inhibition. Discussion and conclusions are provided in sections 4 and 5, respectively.

## 2 Methods

This section describes our simulation framework (summarized in Figure 1), including a multi-fidelity modeling approach (§2.1) that significantly accelerates simulating a coagulation cascade with 32-species reaction kinetics for multiple levels of factor XI/XII inhibition (§2.2). Medical imaging (§2.3), 4D patient-specific mesh generation (§2.4), computational fluid dynamics analyses (§2.5), and verification methods (§2.6) are also described.

**Figure 1:**
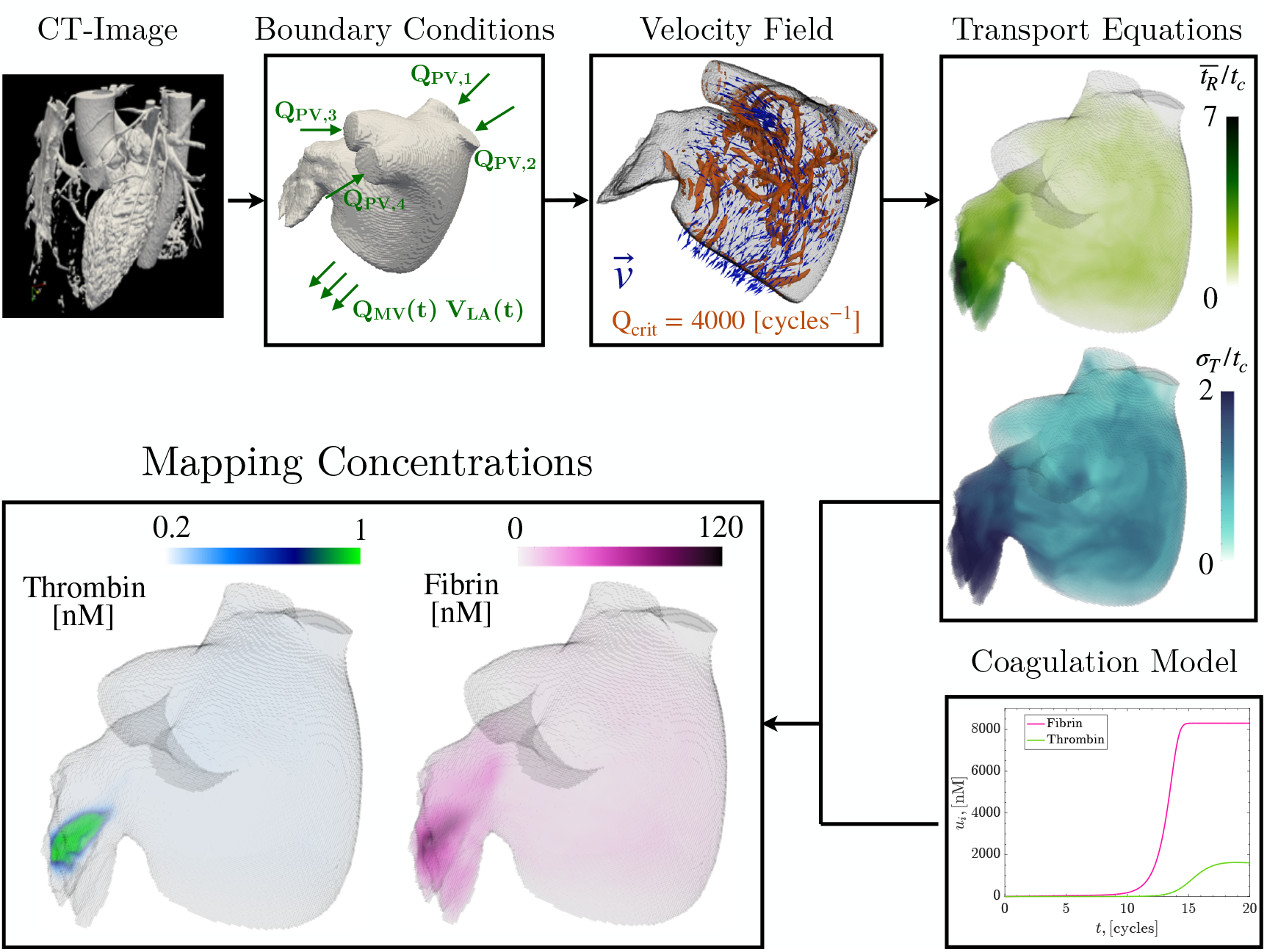
Workflow for Multi-fidelity approach of the coagulation cascade in patient-specific data: The LA wall motion is obtained from CT imaging. Subsequently, the total flow rate through the PVs (*Q*_*PV*_) is calculated from mass conservation in the LA volume and evenly distributed through each PV (*Q*_*PV,i*_). The velocity field 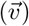, residence time 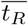, and its higher order moments (e.g., 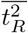) are computed by solving the Navier-Stokes equations for incompressible flow using computational fluid dynamics and transport equations (3–4). A 32-species ODE coagulation model is solved for different levels of factor XI/XII inhibition, and each species’ spatial concentration field is mapped using multi-fidelity (MuFi) modeling (eqs. 6–7).

### 2.1 High-Fidelity (HiFi) and Multi-Fidelity (MuFi) Models of the Coagulation Cascade Under Flow

Considering blood as a continuum flowing with velocity 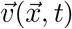, the evolution of the concentration of coagulation species is modeled by a system of reaction-advection-diffusion equations:

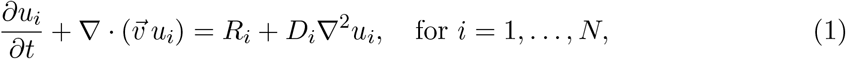

where 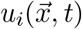 for *i* = 1 … *N* are concentration fields of the involved *N* species. The terms *R*_*i*_(*u*_1_, *u*_2_, …, *u*_*N*_) denote the reaction rates from chemical kinetics, and *D*_*i*_ stand for their diffusivity coefficients. We refer to this system of *N* partial differential equations (PDEs) as the high-fidelity (HiFi) model.

We can non-dimensionalize equations (1) using the flow velocity scale *U*_*c*_ and vessel length scale *L*_*c*_:

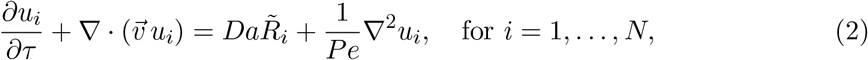

where *τ* = *tU*_*c*_*/L*_*c*_ is a dimensionless time variable, 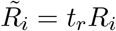 is a dimensionless reaction rate normalized with the characteristic time of the coagulation cascade *t*_*r*_, the Damköhler number *Da* = *L*_*c*_*/*(*t*_*r*_*U*_*c*_) measures the relative importance of reaction kinetics and convective terms, and the Péclet number *Pe* = *U*_*c*_*L*_*c*_*/D*_*i*_ measures the relative importance of convection over diffusion. Given knowledge of 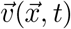, this HiFi model can be solved with appropriate initial and boundary conditions for *u*_*i*_. Dirichlet boundary conditions are enforced at the flow inlets (i.e., on the pulmonary veins for simulations of the LA flow) as *u*_*i*_ = *u*_*i*,0_, while homogeneous Neumann boundary conditions (∂*u*_*i*_*/*∂*n* = 0) are applied at solid surfaces and flow outlets.

Using typical values corresponding to the left atrium and the reaction rate and diffusivity of coagulation cascade species (i.e., *U*_*c*_ ∼ 10 cm/s, *L*_*c*_ ∼ 1 cm, *t*_*r*_ ∼ 10^2^ s, *D*_*i*_ ∼ 10^−6^ cm^2^/s) yields *Da* ∼ 10^−3^ and 1*/P e* ∼ 10^−7^. With a cardiac cycle period of *t*_*c*_ = 1s, equivalent to *t*_*c*_ = 10*L*_*c*_*/U*_*c*_, solving this PDE system implies discretizing the domain into extremely fine grids due to the very large Péclet number, and running it for tens of cardiac cycles due to the Damköhler number of the reaction. Additionally, a complete description of the coagulation cascade typically involves dozens of species (*N* ∼ 50), leading to a large number of PDEs. Furthermore, many practical applications require multiple simulations sweeping over one or more parameters (i.e., initial and/or inlet concentrations of blood clotting factors, kinetic reaction constants, etc.), significantly increasing the compute time.

To alleviate this burden, we employed the Multi-Fidelity (MuFi) approach proposed by Guerrero-Hurtado et al. [2023], where the *N* reaction-advection-diffusion equations for the species concentration are transformed into a set of ODEs using the blood residence time 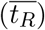 as the independent variable. Therefore, MuFi method involves integrating a single PDE for 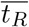, solving a system of *N* ODEs for the coagulatory species, and then mapping the concentration fields as a function of the residence time, 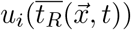.

This transformation is exact in the limit of zero diffusivity. However, with small but finite diffusivity, the approach can be expanded using a Taylor expansion in terms of the statistical moments of residence time, such as 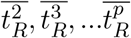. This expansion allows for deriving customizable MuFi models that optimize the trade-off between the computational cost and the order of the expansion. This work employed MuFi models of up to third order involving the evolution equations

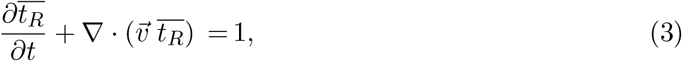

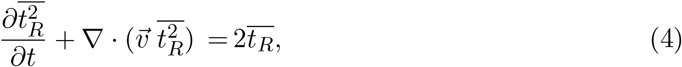

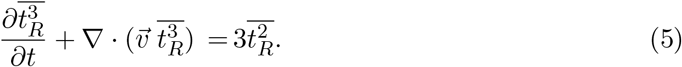

After solving these 3 PDEs, one can map the chemical species concentrations using the Taylor expansions

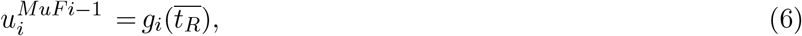

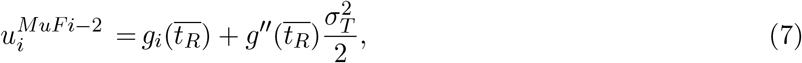

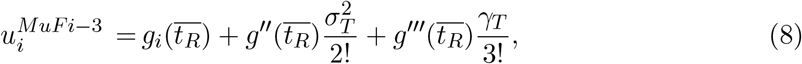

where 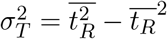 and 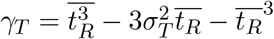 are the second- and third-order moments of the residence time centered in the mean. In these equations, *g*_*i*_, 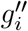, and 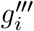 represent the solution and time derivatives of the concentration of the chemical species *i*, determined by solving a system of N ODEs governing the dynamics of a well-mixed fluid volume with homogeneous initial conditions 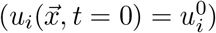:

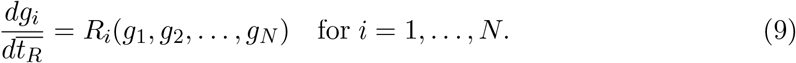

We refer to eq. (9) as the no-flow reaction model. For the results presented in section 3, the no-flow reaction model was integrated in time using an explicit, low-storage, 3-stage Runge-Kutta scheme. Once residence time and its higher-order moments are computed, the MuFi allows for evaluating the coagulation cascade under multiple conditions by integrating the no-flow ODE system at almost negligible cost.

### 2.2 Factor XI/XII anticoagulant simulations: reaction kinetics and clotting metrics

We implemented a 32-species coagulation system with the reaction kinetics described by Zhu [2007] in our MuFi model of factor XI/XII anticoagulants. This system is an adaptation of the system proposed by Kogan et al. [2001], which includes detailed activation of factors XI and XII, and the reactions leading to the subsequent activation of factor X. We defined a prothrombotic initial condition with the concentrations reported in Table 1, setting the concentrations of all active factors to zero except for thrombin (IIa) and factor XIIa.

**Table 1:** Nominal Initial Concentrations.

| Factor | Concentration [$\mu\text{M}$] | Reference |
| --- | --- | --- |
| PK | 0.58 | Saito et al. [1978] |
| XIIa | $2.3 \cdot 10^{-5}$ | Zhu [2007] |
| C1 | 1.7 | Harpel [1987] |
| PAI | $4.6 \cdot 10^{-4}$ | Kruithof et al. [1988] |
| $\alpha_2\text{M}$ | 3.5 | Harpel [1987] |
| ATIII | 3.4 | Collen et al. [1977] |
| XII | 0.6 | Madsen et al. [2013] |
| XI | 0.06 | Gailani and Broze Jr [1991] |
| $\alpha_2\text{AP}$ | 0.9 | Harpel [1987] |
| IX | 0.18 | Komiyama et al. [1990] |
| X | 0.34 | Tormoen et al. [2013] |
| TFPI | $2.5 \cdot 10^{-3}$ | Novotny et al. [1989] |
| $\alpha_1\text{AT}$ | 24.5 | Harpel [1987] |
| IIa | $2 \cdot 10^{-4}$ | - |
| II | 1.8 | Monroe et al. [2020] |
| V | 0.042 | Tracy et al. [1982] |
| TM | $2.2 \cdot 10^{-4}$ | Aso et al. [2000] |
| PC | 0.064 | Vaziri et al. [1988] |
| VIII | $1.4 \cdot 10^{-3}$ | Butenas et al. [2010] |
| I | 8.3 | Ratnoff and Menzie [1951] |

We modeled either factor XI or factor XII anticoagulant treatment by inhibiting each of these factors’ initial concentration. We defined the inhibition level of factor *i* as

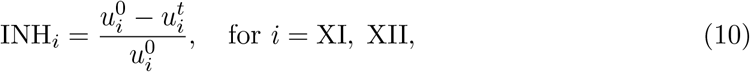

where 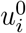 is the nominal concentration (see Table 1) and 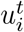 is the inhibited target concen-tration. For the simulations presented in section 3.4 we consider 9 inhibition levels for each factor, i.e., INH_*i*_ = [0.25, 0.50, 0.70, 0.75, 0.80, 0.85, 0.90, 0.95, 0.975].

We employed two metrics to evaluate the resulting coagulation cascade in each patient-specific simulation. Similar to previous works [Nicoud, 2023, Danforth et al., 2012], we defined the coagulation time *t*_*co*_ as the time at which thrombin concentration reaches a threshold value 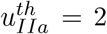 nM anywhere inside the LAA. Analogous to the clotting times defined in clinical tests (i.e., PT, aPTT, TT, etc), *t*_*co*_ measures the time required to activate the coagulation cascade in flowing blood. Additionally we defined the coagulating volume as

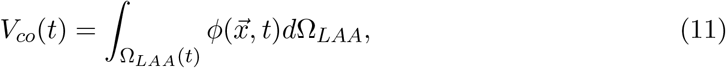

which measures the volume within the LAA where the coagulation cascade is activated. In this equation, Ω_*LAA*_(*t*) indicates the set of points 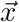 belonging to the LAA, 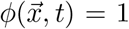 at points where 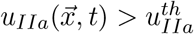, and 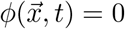 elsewhere.

### 2.3 CT Imaging

We studied a group of *N* = 13 patients, selected to sample a wide range of anatomical and functional characteristics relevant to atrial fibrillation and thrombosis. Subjects 1-3 were enrolled at the National Institutes of Health (NIH) in Bethesda, Maryland (N = 3). Subjects 4-11 were enrolled at the University of California San Diego (UCSD), CA, United States (N = 8); Subjects 12 and 13 were enrolled at Hospital General Universitario Gregorio Marañón (HGUGM), Madrid, Spain. The studies received approval from the Institutional Review Board at all three centers.

Each of the 13 study patients underwent 3D, time-resolved computed tomography scans (4D-CT) to segment the LA anatomy. The voxel dimension ranged from 0.32 mm to 0.62 mm in the x-y plane and from 0.45 mm to 1 mm in the z direction. Eight of these subjects were included in our previous studies [Garcia-Villalba et al., 2021, Gonzalo et al., 2022, Durán et al., 2023], and five new patients were added. Time-resolved imaging data were obtained at regular intervals across the cardiac cycle, spanning from 5% to 10% of the R-R interval.

### 2.4 4D Patient-Specific LA Meshing

The LA computational meshes were generated in four steps using ITK-SNAP [Yushkevich et al., 2006] and custom MATLAB scripts. Initially, the 3D LA anatomy was segmented from CT images, identifying key landmarks such as the pulmonary vein (PV) inlets, mitral annulus, and left atrial appendage (LAA). Then a triangular mesh was created for each LA segmentation with the resolution of the CFD solver [Fang, 2009]. These meshes were registered across the cardiac cycle to ensure coherence in vertex and centroid positions [Myronenko and Song, 2010]. Subsequently, these point positions were converted into Fourier temporal series, providing the interpolated boundary conditions to the CFD solver at time points between the time frames of the 4D CT sequence. Further details on image acquisition, reconstruction, and mesh generation can be found in Garcia-Villalba et al. [2021].

### 2.5 Computational Fluid Dynamics

We adapted the CFD code TUCANGPU [Guerrero-Hurtado et al., 2024] to solve the Navier-Stokes equations for incompressible flow

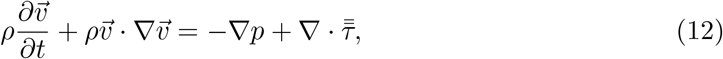

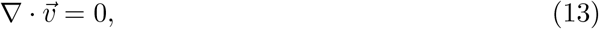

where 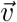 and *p* are the velocity and pressure fields, *ρ* the fluid density, and 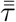 the viscous stress tensor, using patient-specific LA meshes. We used a residence-time-activated Carreau-Yasuda model [Gonzalo et al., 2022] to represent the thixotropic, shear-thinning rheology of blood arising from formation and rupture of RBC aggregates [Robertson et al., 2008]. This model provides a non-Newtonian constitutive relation between blood viscosity *ν* and shear rate *S* that depends on 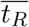 as

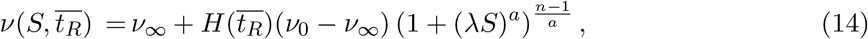

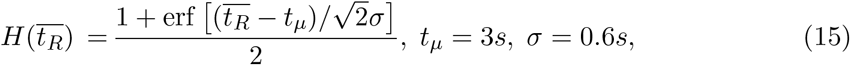

where *t*_*µ*_ are timescales associated to RBC aggregation, and *λ* = 8.2, *a* = 24.32, *n* = 0.37, *ν*_0_ = 16*ν*_∞_ with *ν*_∞_ = 0.04cm^2^*/*s are the Carreau-Yasuda model constants. These constants depend on the hematocrit (*Hct*) of the patient, and the values chosen for our simulations correspond to *Hct* = 43.5, which falls within the physiological range [Walker et al., 1990]. Figure 2 illustrates the impact of residence time 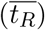 and shear (*S*) on the viscosity (*ν*), showing that non-Newtonian effects are negligible for 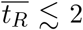 s, gradually increasing for 2 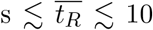 s. For 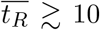 s the kinematic viscosity *ν* provided by the residence-time-activated Carreau-Yasuda model becomes indistinguishable from its classic version.

**Figure 2:**
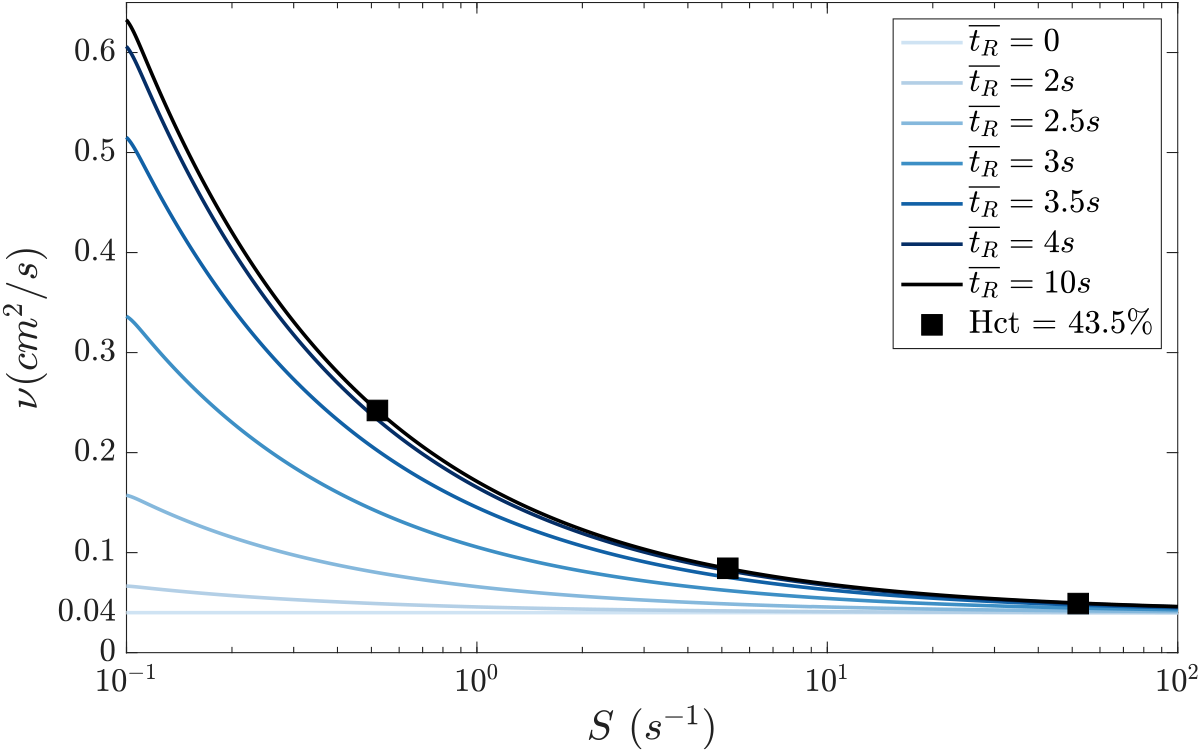
Residence-Time-Activated Carreau–Yasuda model: Non-Newtonian constitutive laws for the kinematic viscosity as a function of shear rate (*S*) and residence time 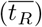, denoted as 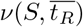, for the residence-time-activated Carreau–Yasuda model. The black squares represent data from Chien et al. [1966] study for the respective hematocrit (Hct) value.

Each patient-specific LA simulation was run for 20 cardiac cycles with a fixed time step Δ*t*, chosen to ensure a Courant-Friderichs-Lewy (CFL) number below 0.3 throughout the complete run. The fluid domain was discretized using a staggered Cartesian grid with a constant spacing of Δ*x* = 0.051 mm and the spatial derivatives were approximated with centered second-order finite differences. The segmented LA geometry was embedded within a 13-cm cubic domain with periodic boundary conditions. The LA surface motion, derived from patient-specific 4D CT images, was prescribed throughout the cardiac cycle and influenced flow via the no-slip boundary condition, which was enforced using the immersed boundary method (IBM) proposed by Uhlmann [2005].

Inflow boundary conditions were imposed assuming equal flow rate through each pulmonary vein (PV), denoted as *Q*_*i*_, for *i* = 1…4. Specifically,

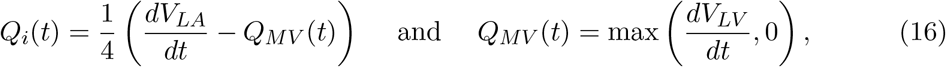

where *V*_*LA*_ represents the time-dependent volume of the LA, *Q*_*MV*_ denotes the flow rate through the mitral valve, and *V*_*LV*_ is the left ventricle (LV) volume obtained from the CT image. To enforce the velocity *v*_*i*_ at each PV, a non-cylindrical buffer region of Lagrangian points was extended upstream by twelve grid planes. This inlet velocity was imposed within the flow domain using a volumetric force, akin to the IBM force.

Boundary conditions at the mitral annulus outlet were applied to the plane section at the downstream end of the atrial segmentation, which dynamically moved within the cubic simulation domain as the LA walls deformed. When the mitral valve was closed, mesh points in that section were treated as a standard no-slip boundary, identical to the rest of the atrial wall. Conversely, when the mitral valve was open (i.e., *Q*_*MV*_ > 0), no boundary condition (i.e., no IBM forcing) was imposed on these mesh points.

The transport equations for the residence time and higher order moments (eqs. 3, 4, and 5) were solved together with the Navier-Stokes equations in the TUCANGPU CFD solver. To address the lack of diffusion on eqs. (3), (4), and (5), we followed our previous work [Garcia-Villalba et al., 2021, Gonzalo et al., 2022, Durán et al., 2023], and employed a third-order weighted essentially non-oscillatory (WENO) scheme [Jiang and Shu, 1996] to compute the non-linear terms. This scheme prevents spurious oscillations in the numerical solutions while minimizing the overall artificial diffusivity. For the MuFi approach, the concentration field 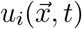 was mapped from the residence time and higher order moments using eqs. (6), (7) or (8), with the values of *g*_*i*_, 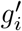 and 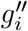 obtained from the no-flow ODE systems described in section 2.2.

### 2.6 Verification of MuFi modeling in 3D patient-specific anatomies

To verify the MuFi models, we solved the HiFi reaction-advection-diffusion equations numerically (2) in 3D patient-specific LA anatomies and compared the results of the MuFi and HiFi models. We considered models of first (MuFi-1), second (MuFi-2), and third order (MuFi-3) to evaluate which MuFi order was most efficient at balancing computational cost with accuracy.

Given the high cost of running the HiFi system, we made several arrangements to reduce compute time. Similar to our previous work [Guerrero-Hurtado et al., 2023], we used the 9-species system of Zarnitsina et al. [2001] instead of the 32-species of Zhu [2007] with the reaction rates and initial conditions described in Appendix A1. Instead of performing the verification analysis for all patients, we selected two subjects representative of normal and impaired atrial function.

The numerical discretization of the HiFi system of PDEs was similar to that of the PDEs governing the residence time and included a WENO scheme for the non-linear terms. However, to further reduce compute time, the HiFi system was not integrated together with the flow in the CFD solver. Instead we solved the incompressible Navier-Stokes equations (eq. 12) over 10 cycles to ensure a quasi time-periodic flow. Then, we phase averaged the last 5 cycles, and stored 40 3D velocity fields, i.e., one field every 500 time steps. Subsequently, we solved the HiFi PDE system interpolating the 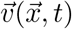 linearly in time. To ensure an unbiased comparison between the HiFi and MuFi models, the residence time and higherorder moments used in the verification study were obtained solving transport equations (3), (4) and (5) with linearly interpolated velocity fields in time. For all other results reported in this manuscript, we used the full-resolution residence time fields integrated concurrently with the flow in the CFD solver, as described in §2.5 above.

The global relative error of the MuFi models inside the LAA was quantified as

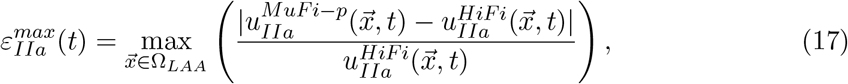

where Ω_*LAA*_ represents the volume of the LAA at each time step of the simulation. We focused on the LAA because this is the site of maximum thrombin concentration and most likely site of atrial thrombosis.

## 3 Results

This section begins with a description of our subjects (§3.1) and verification study of the Multi-Fidelity (MuFi) approach in realistic left atrial (LA) flows (§3.2). Subsequently, the spatio-temporal evolution of the coagulation cascade in 13 patient-specific LA flows is analyzed (§3.3). Finally, §3.4 evaluates the effectiveness of two anticoagulant therapies targeting the intrinsic pathway via factor XI/XII inhibition.

### 3.1 Patient characteristics

Table 2 summarizes patient demographics and atrial anatomical and functional characteristics relevant to fibrillation and thrombosis. The median age of the group was 65 years and 54% were female. Subjects 1-3, 12, and 13 were imaged in sinus rhythm, showing no evidence of LAA thrombus and demonstrating normal LA function, as indicated by chamber volumes and PV flows derived from CT images. Conversely, subjects 4-11 exhibited an enlarged LA with impaired global function. Subjects 5, 6, and 11 had an LAA thrombus, which was digitally removed from the segmentations before running the CFD simulations.

**Table 2:** Anatomical and functional parameters of the LA and the LAA. The mean volume values represent time-averaged volumes. The emptying fraction for LA and LAA are defined as *EF*_*LA*_ = (max(*V*_*LA*_) − min(*V*_*LA*_)*/* max(*V*_*LA*_) and *EF*_*LAA*_ = (max(*V*_*LAA*_) − min(*V*_*LAA*_)*/* max(*V*_*LAA*_), respectively. TIA stands for Transient Ischemic Attacks.

| Subject | 1 | 2 | 3 | 4 | 5 | 6 | 7 | 8 | 9 | 10 | 11 | 12 | 13 |
| --- | --- | --- | --- | --- | --- | --- | --- | --- | --- | --- | --- | --- | --- |
| LAA thrombus | No | No | No | TIA | Yes | Yes | No | No | No | No | Yes | No | No |
| Persistent AF | No | No | No | No | Yes | Yes | Yes | Yes | Yes | Yes | Yes | No | No |
| Sinus rhythm | Yes | Yes | Yes | Yes | No | No | No | No | No | Yes | No | Yes | Yes |
| Mean LA vol, (ml) | 86.6 | 70.1 | 115 | 145 | 157 | 180 | 229 | 177 | 193 | 132 | 150 | 84 | 96 |
| Max LA vol, (ml) | 108 | 91.2 | 145 | 155 | 165 | 205 | 247 | 194 | 208 | 157 | 160 | 108 | 121 |
| Min LA vol, (ml) | 59.6 | 49.0 | 87.2 | 119 | 150 | 157 | 216 | 169 | 183 | 116 | 139 | 60.3 | 68.6 |
| Mean LAA vol, (ml) | 6.94 | 4.85 | 14.3 | 10.7 | 15.5 | 22.0 | 5.51 | 6.17 | 14.1 | 14.1 | 15.7 | 3.13 | 3.64 |
| Max LAA vol, (ml) | 8.97 | 6.28 | 17.9 | 11.6 | 17.4 | 24.7 | 6.58 | 7.41 | 15.47 | 17.0 | 18.5 | 4.23 | 4.65 |
| Min LAA vol, (ml) | 4.32 | 3.14 | 10.2 | 9.10 | 13.8 | 19.8 | 5.01 | 5.17 | 13.4 | 12.5 | 13.16 | 1.45 | 2.24 |
| $EF_{LA}$ | 0.45 | 0.46 | 0.4 | 0.23 | 0.09 | 0.23 | 0.123 | 0.127 | 0.12 | 0.26 | 0.132 | 0.44 | 0.43 |
| $EF_{LAA}$ | 0.52 | 0.50 | 0.43 | 0.2 | 0.21 | 0.22 | 0.24 | 0.30 | 0.13 | 0.26 | 0.28 | 0.66 | 0.52 |
| Anticoagulants | No | No | No | No | Yes | Yes | Yes | Yes | Yes | Yes | Yes | No | No |

Subject 4 did not have an LAA thrombus but had a history of transient ischemic attacks (TIAs). Based on these data, we identified subjects 4-11 as AF positive and subjects 5-6 and 11 as clot-positive. Figure 3 shows the segmented LA anatomies of the 13 patients at the onset of the R-R interval, including inlet (PVs) and outlet (mitral annulus) sections. The color assigned to each case is based on its clinical diagnosis: red for clot-positive cases, blue for clot-negative cases that are AF positive, and green for AF negative cases.

**Figure 3:**
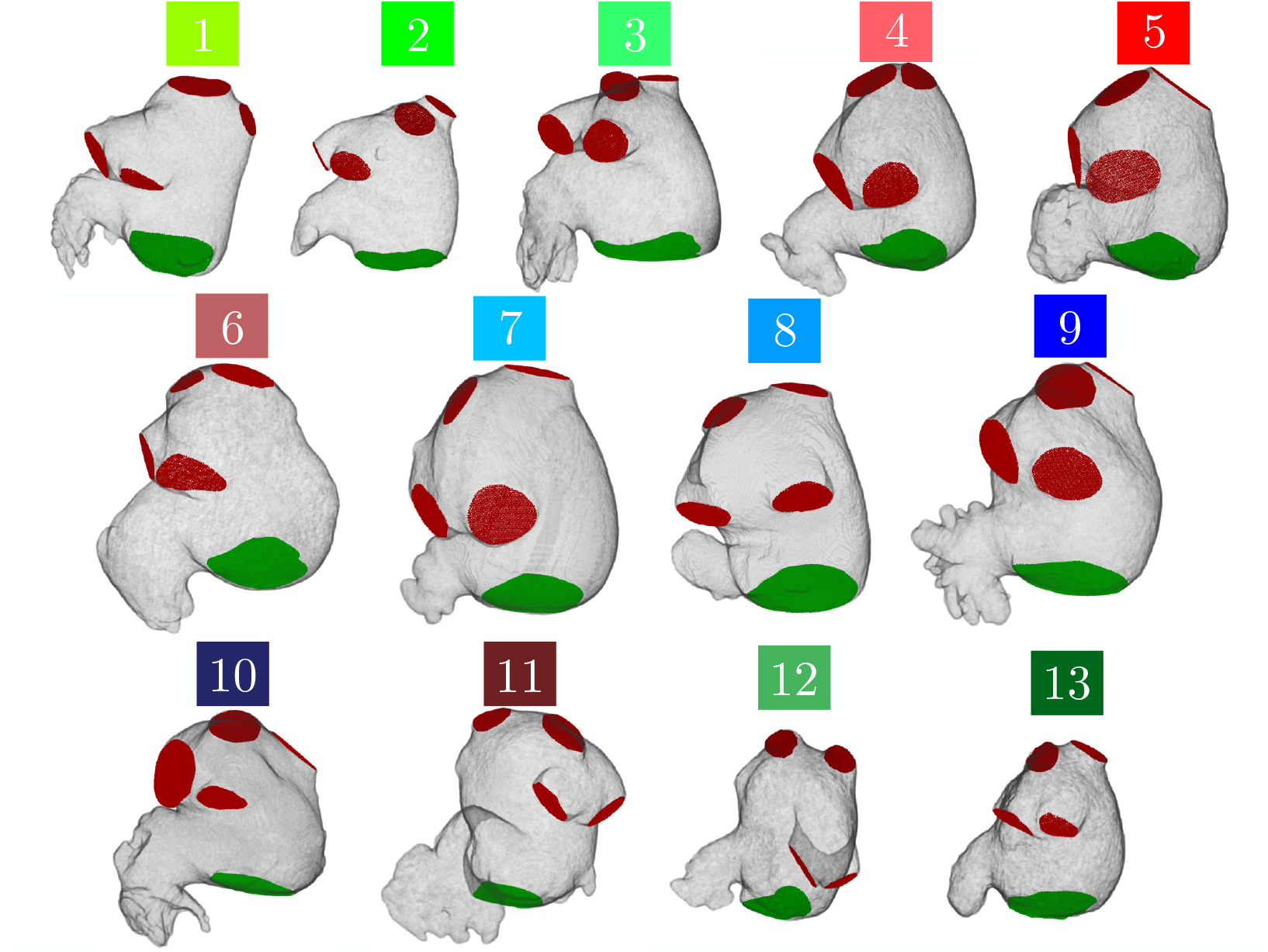
Anatomical Configuration of Patient-Specific Left Atrium Subjects: Three-dimensional mesh derived from Computerized Tomography (CT) scans depicting the left atrium walls and pulmonary veins (PVs highlighted in red) as well as the mitral valve outlet surfaces (in green). These images capture a moment at the start of the R-R interval.

### 3.2 Comparison of Multi-Fidelity and High-Fidelity coagulation models

Multi-fidelity (MuFi) modeling of reaction-advection-diffusion processes in the low-diffusivity limit has shown promise as a means to significantly accelerate the simulation of blood coagulation under flow [Guerrero-Hurtado et al., 2023]. However, this approach has not been verified yet in realistic 3D cardiovascular geometries. This section compares HiFi and MuFi simulations of a 9-species coagulation system for 3D patient-specific LA flows corresponding to one LAA-thrombus-negative (case 2) and one LAA-thrombus-positive individual (case 6).

Figure 4 depicts the spatial distributions of the thrombin concentration, *u*_*IIa*_, for the HiFi model and the MuFi-1, MuFi-2, and MuFi-3 models. The data are represented at early atrial diastole of the 20th simulation cycle. Consonant with the HiFi model, the three MuFi models captured that thrombin concentration peaks in the LAA and predicted significantly higher *u*_*IIa*_ in the thrombus-positive patient (Figure 4A-D) than in the thrombus-negative patient (Figure 4E-H). Nevertheless, MuFi-1 tended to underestimate the peak values of *u*_*IIa*_, whereas MuFi-2 and MuFi-3 produced *u*_*IIa*_ distributions in closer agreement with the HiFi results.

**Figure 4:**
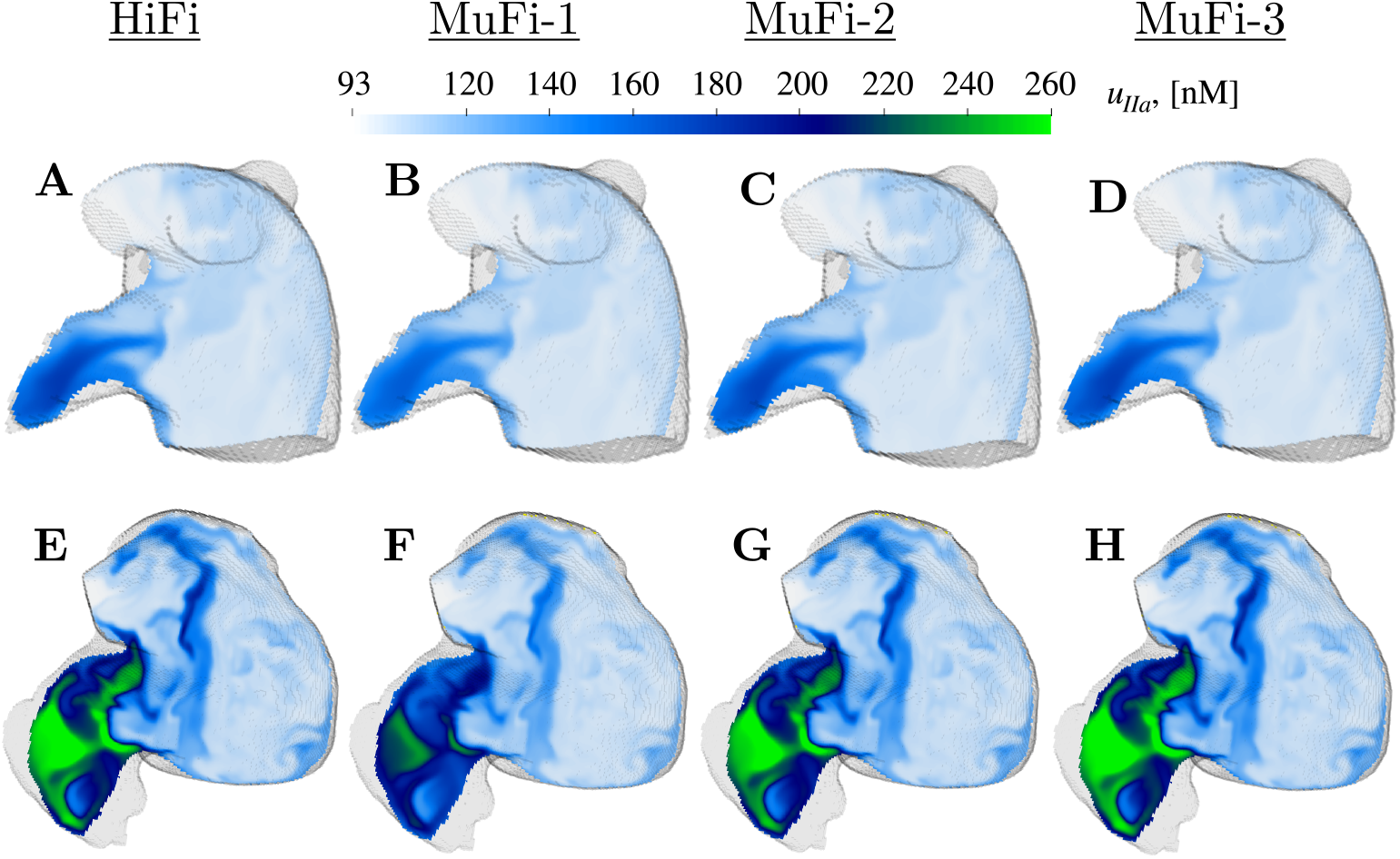
Spatial distribution of thrombin concentration *u*_*IIa*_ on oblique plane sections: Case 2 (top) and case 6 (bottom) at *t/t*_*c*_ = 20. HiFi (A,E), Mufi-1 (B,F), MuFi-2 (C,G), MuFi-3 (D,H)

To conduct a more detailed evaluation of the MuFi models, we compared the temporal evolutions of *u*_*IIa*_ predicted at the ostium center for the same two patients (Figure 5). In these plots, the 9-species no-flow reaction model was also included for reference (solid line). Both the HiFi and all MuFi models departed from the no-flow model quickly after one cardiac cycle, revealing that no-flow models severely overestimate thrombin concentrations for timescales relevant to the coagulation process. Thrombin concentration in the HiFi and MuFi models experienced oscillations with a slowly growing envelope due to the cyclic inflow of “fresh” low-*u*_*IIa*_ blood. In the more contractile atrium (case 2, Figure 5A), this fluid exchange was more vigorous and, consequently, the intra-cycle peak value of *u*_*IIa*_ stopped growing at *t* ≈ 8*t*_*c*_. On the other hand, the envelope of *u*_*IIa*_ was still growing slowly at *t* = 20*t*_*c*_ for the less contractile atrium (case 6, Figure 5B).

**Figure 5:**
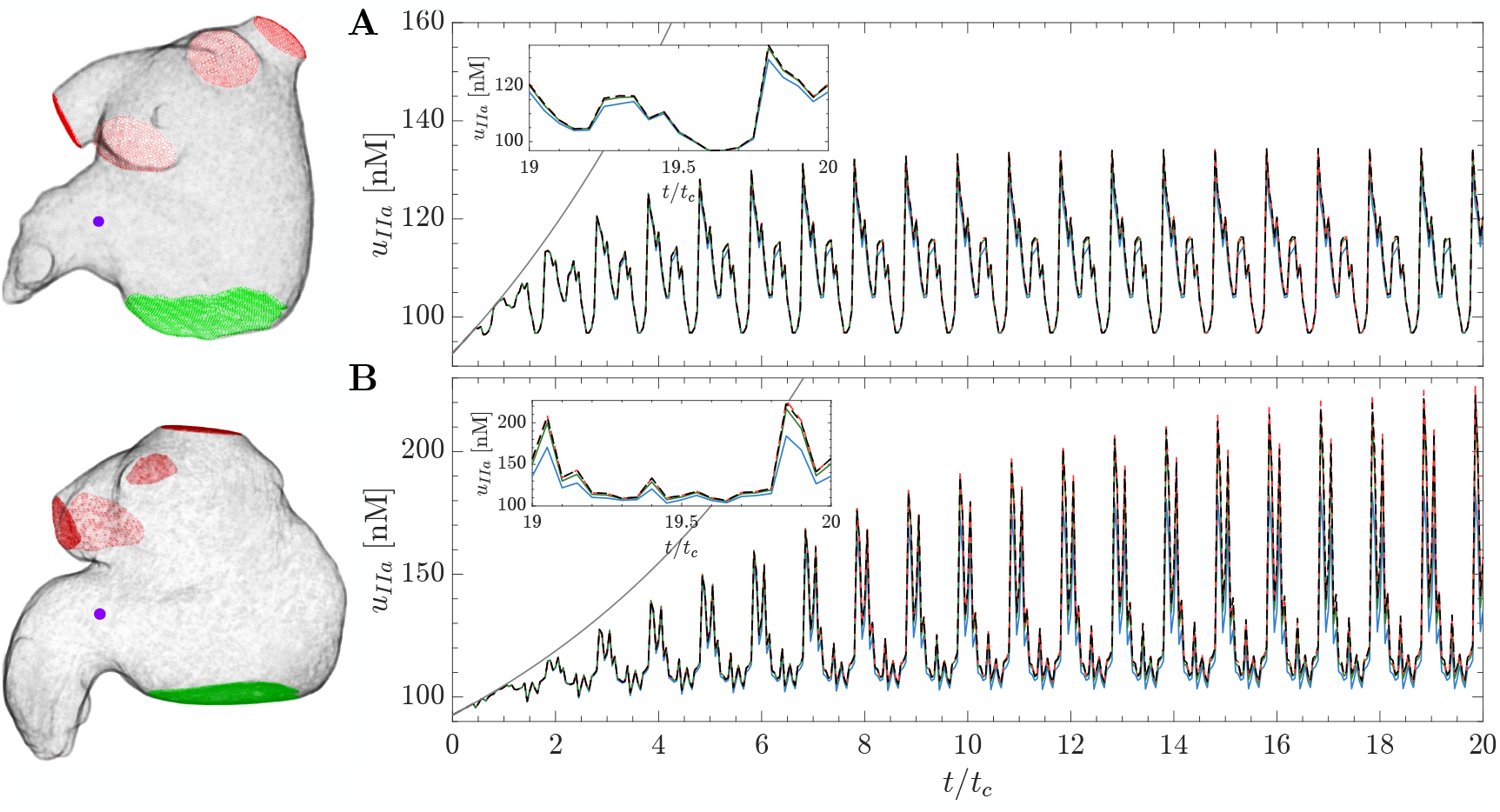
Time series of thrombin concentration *u*_*IIa*_: Case 2 (A) and Case 6 (B). Each line corresponds to a different model: HiFi 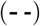, MuFi-1 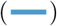, MuFi-2 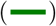 and MuFi-3 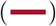. The locations are considered in the near region of the ostium plane and indicated with 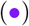. For reference, the solution of the 9-species no-flow reaction model (eq. 9) is also included 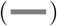.

Overall, all the MuFi models captured the temporal dynamics of *u*_*IIa*_ regardless of their order. However, consistent with the results in Figure 4, MuFi-1 tended to underestimate the peak values of *u*_*IIa*_ after integrating the models for 20 cardiac cycles. These differences were more significant in the patient exhibiting higher thrombin concentrations (case 6, Figure 5B). The MuFi-2 model showed good agreement with the HiFi solution and the MuFi-3 model further seemed to capture the concentration peaks slightly better.

Figure 6 displays the time evolution of the global relative error of the MuFi models, 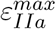 defined in eq. (17), for the three MuFi models and two patients discussed above. In both patients, MuFi-1 exhibited markedly larger errors than the other MuFi models at all times, reaching *ε*^*max*^ ∼ 10% in case 2 and 25% in case 6 by the 20th cycle. The MuFi-2 and MuFi-3 errors differed less from each other and were smaller than 0.3% for the first 5 cardiac cycles. For *t* ≳ 5*t*_*c*_, the MuFi-2 error grew faster with time, reaching *ε*^*max*^ ∼ 3% and 8% for cases 2 and 6 at *t* = 20*t*_*c*_, while the MuFi-3 model had *ε*^*max*^ ∼ 1% and 2%.

**Figure 6:**
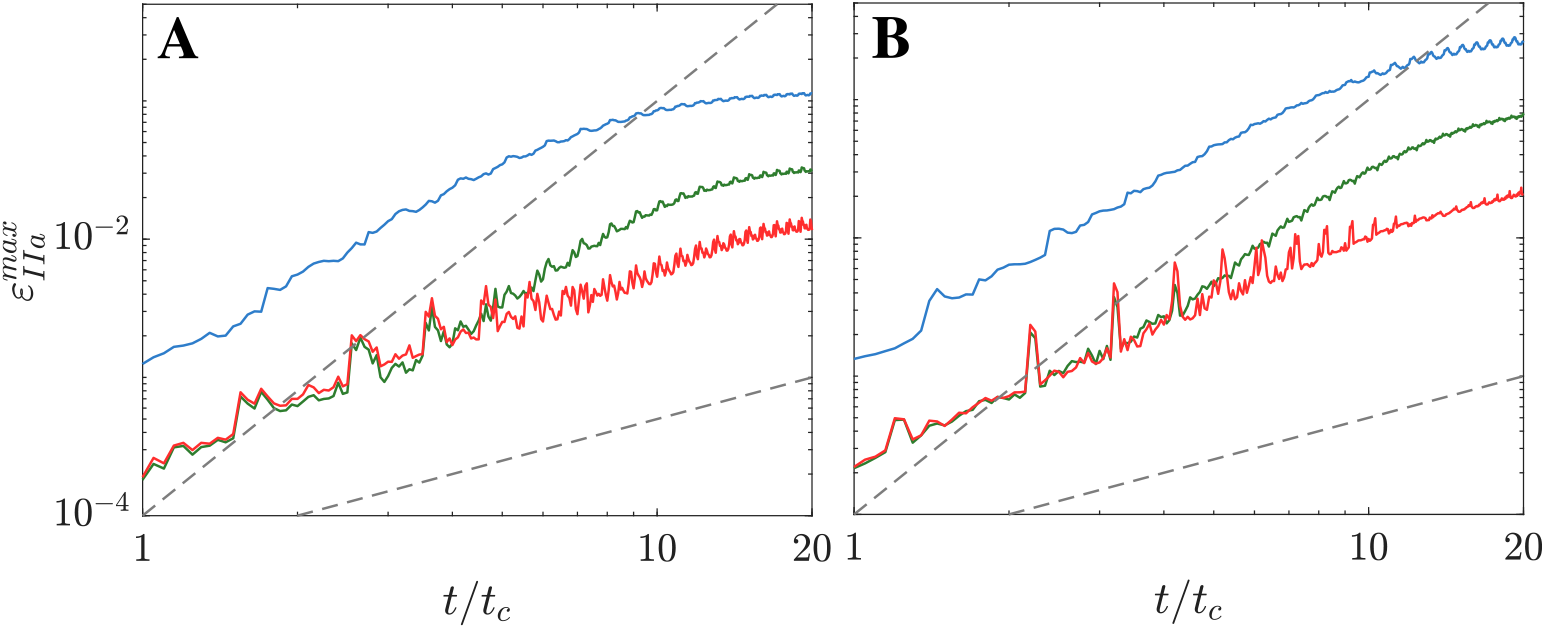
Maximum relative error in the LAA: Case 2 (A) and Case 6 (B). Each line corresponds to a different model: MuFi-1 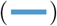, MuFi-2 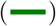 and MuFi-3 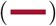. Dashed-dot lines correspond to 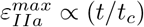 and 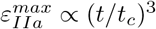.

In terms of computational cost, solving 20 cardiac cycles of the 9-equation coagulation system using the HiFi approach took approximately 320 minutes using the Python version with Numba CUDA implementation, as detailed in Guerrero-Hurtado and Flores [2023], on a GPU A100 with 80 GB of RAM, 6912 CUDA cores, and 432 tensor cores. Conversely, on the same hardware, the MuFi-1 approach took approximately 35 minutes, while the MuFi-2 and MuFi-3 approaches required roughly 45 minutes and 72 minutes, respectively. Based on these results, the subsequent analyses presented in this manuscript utilized the MuFi-2 model in an effort to balance accuracy and computational efficiency.

### 3.3 Patient-Specific Models of LA Coagulation Cascade Initiation

This section examines the progression of thrombin concentration in 13 3D patient-specific LA models by applying a MuFi-2 model with 32-species kinetics. The analysis uses the nominal initial concentrations outlined in Table 1 and temporal integration is performed over 20 cardiac cycles. The velocity fields, 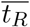, and 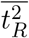 are all obtained by the CFD solver described in §2.5, therefore eliminating the need for temporal interpolation.

Figure (7A) illustrates the temporal evolution of the maximum thrombin concentration within the LAA normalized with the threshold concentration value 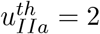 nM, and Figure (7B) displays the temporal evolution of the normalized LAA coagulating volume, *V*_*co*_. As before, the results from the no-flow ODE model are included for reference in Figures 7A-B (dashed line). In this model, the thrombin concentration starts to grow exponentially in the 8-th cardiac cycle, reaching the threshold concentration at *t* ≈ 9.6*t*_*c*_ (Figure 7A). Consequently, the normalized LAA volume is a step function that jumped from zero to one at that time point (Figure 7B).

**Figure 7:**
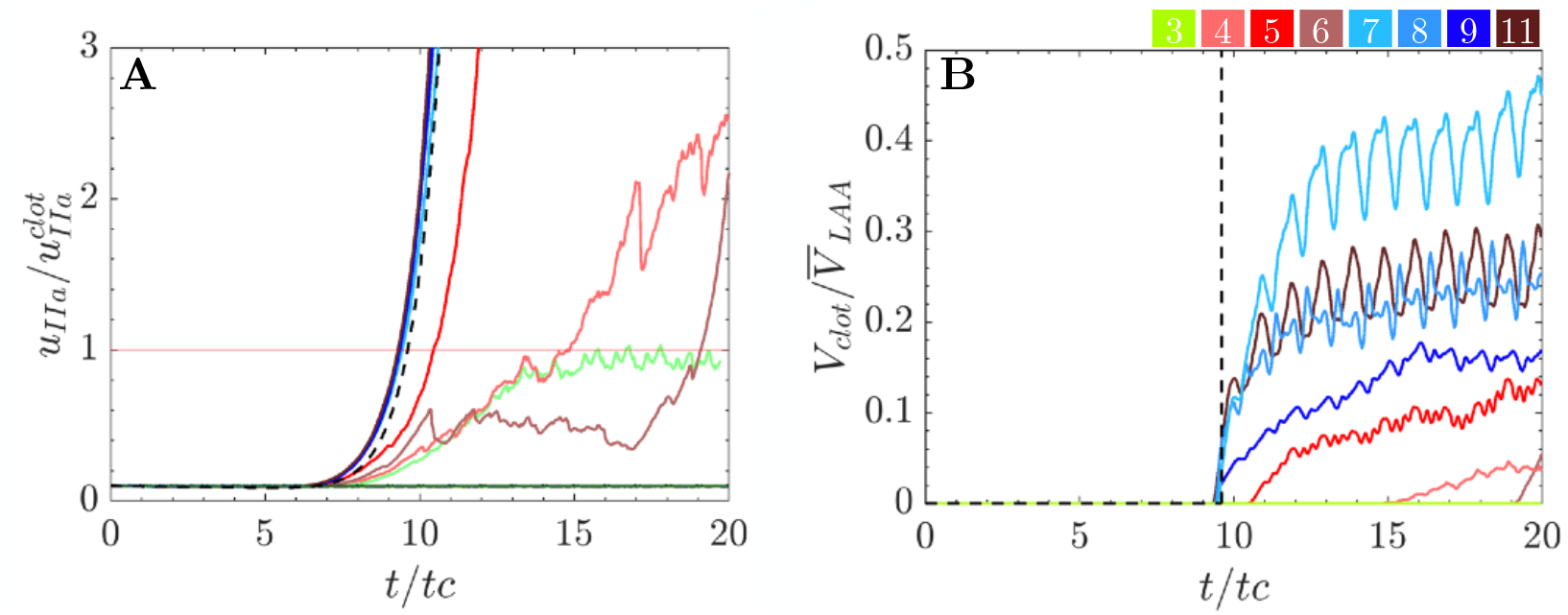
Temporal evolution of thrombin concentration and coagulating volume in the LAA: (A) Time series depicting the maximum thrombin concentration (*u*_*IIa*_) in the LAA of each patient, normalized by the thrombin concentration threshold 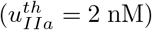. Additionally, the no-flow solution (eq. 9) is provided for reference 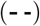. (B) Time series illustrating the coagulating volume within the LAA (*V*_*co*_) for each patient, normalized by the mean LAA volume. Line colors are defined in Figure 3.

Compared to the no-flow results, Figures 7A-B reveal three distinct behaviors in patientspecific LA thrombin dynamics. A first group of *non-coagulating* atria included cases 1, 2, 10, 12, and 13, which exhibited virtually no activation of the coagulation cascade, with nearly constant thrombin values around the initial concentration (horizontal lines in Figure 7A). As a result, all these cases sustained zero normalized coagulating volume over the course of the simulations (Figure 7B). A second group of *moderately coagulating* atria was formed by cases 3, 4 and 6, which experienced more or less intricate patterns of sub-exponential growth and fluctuations in *u*_*IIa*_. The patients in this group had non-zero albeit small values of normalized LAA coagulating volume by *t* = 20*t*_*c*_ (Figure 7B). Finally, a third group of *severely coagulating* atria was formed by cases 5, 7, 8, 9 and 11, which experienced nearly exponential growth with little fluctuations in the maximum thrombin values, similar to the no-flow model (Figure 7A). This result suggests that these cases had recirculating regions inside the LAA that were effectively isolated from the rest of the atrial flow. Accordingly, their normalized LAA coagulating volume profiles resembled a step function with mild intra-cycle oscillations (Figure 7B).

It should be noted that this classification is not overly dependent on the thrombin threshold. For instance, varying this threshold between 1 and 4 nM would only switch case 3 between the non-coagulating and the moderately coagulating groups, whereas all the other cases would not change their classification.

Figure 8 displays the LAA thrombin distribution at *t* = 20*t*_*c*_ in representative cases of the *moderately* and *severely coagulating* groups. The figure does not include *non-coagulating* cases because thrombin was very diluted inside their LAA, rendering the maps non-informative. Also, we used a color scheme with a sharp gradient at the thrombin threshold 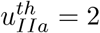 to help visualize each patient’s *V*_*co*_. A noticeable yet moderately sized region of elevated thrombin concentration was observed in the distal LAA of the *moderately coagulating* cases (Figure 8A and B). In one of the cases, the thrombin blob was sub-threshold while in the other case it contained a small area over the threshold. In contrast, the two *severely coagulating* cases had voluminous areas of above-the-threshold thrombin concentration that occupied significant portions of the LAA (Figure 8C and D).

**Figure 8:**
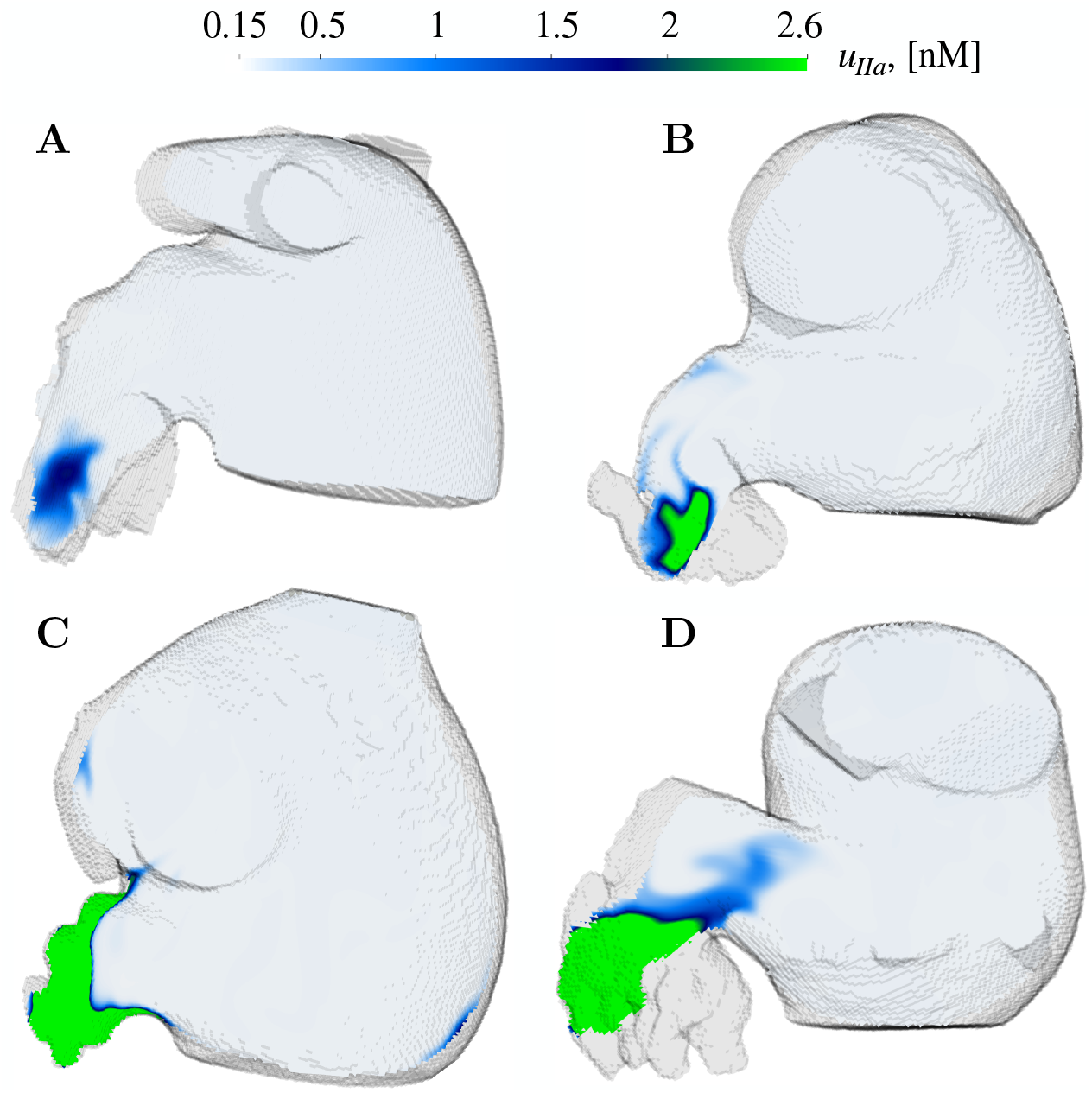
Spatial distribution of thrombin accumulation (*u*_*IIa*_) in oblique plane sections: Thrombin concentration fields (*u*_*IIa*_) in oblique plane sections for *moderately coagulating* cases 3 (A) and 4 (B), and *severely coagulating* cases 7 (C) and 11 (D), after 20 cardiac cycles *t/t*_*c*_ = 20. The color scheme has a sharp jump between dark blue and bright green at *u*_*IIa*_ = 2 nM to facilitate visualizing each patient’s *V*_*co*_.

### 3.4 Digital anticoagulation via factor XI/XII inhibition

We leveraged the computational efficiency of MuFi models to systematically investigate how factor XI/XII inhibition affects LA coagulation under patient-specific flow patterns. Inhibition was modeled by lowering these factors’ initial concentrations from the nominal values shown in Table 1. For each patient, we examined no inhibition and 9 different inhibition levels applied to non-active factors XI or XII, as defined by varying the parameter INH_*i*_ (eq. 10 in §2.2) in the range [0.25, 0.975].

Figure 9 shows the time evolution of the maximum thrombin concentration within the LAA for varying inhibition levels of factor XI. In this figure, we only included data from those patients exhibiting appreciable thrombin concentration after 20 cycles of simulation time, i.e., the *moderately coagulating* and *severely coagulating* groups described in the preceding section. For reference, the results from the flow and no-flow models without inhibition are included in each plot. In all cases shown, the maximum thrombin concentration is normalized by the threshold value 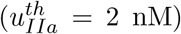. In the *moderately coagulating* group (cases 3, 4, and 6), the raise of thrombin concentration was blunted markedly by moderate inhibitions of factor XI corresponding to INH_*i*_ ≤ 0.5. On the other hand, the *severely coagulating* group was less sensitive to inhibiting factor XI. The patients in this group consistently exceeded the threshold 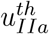 within the 20 simulated cycles for all levels of inhibition considered.

**Figure 9:**
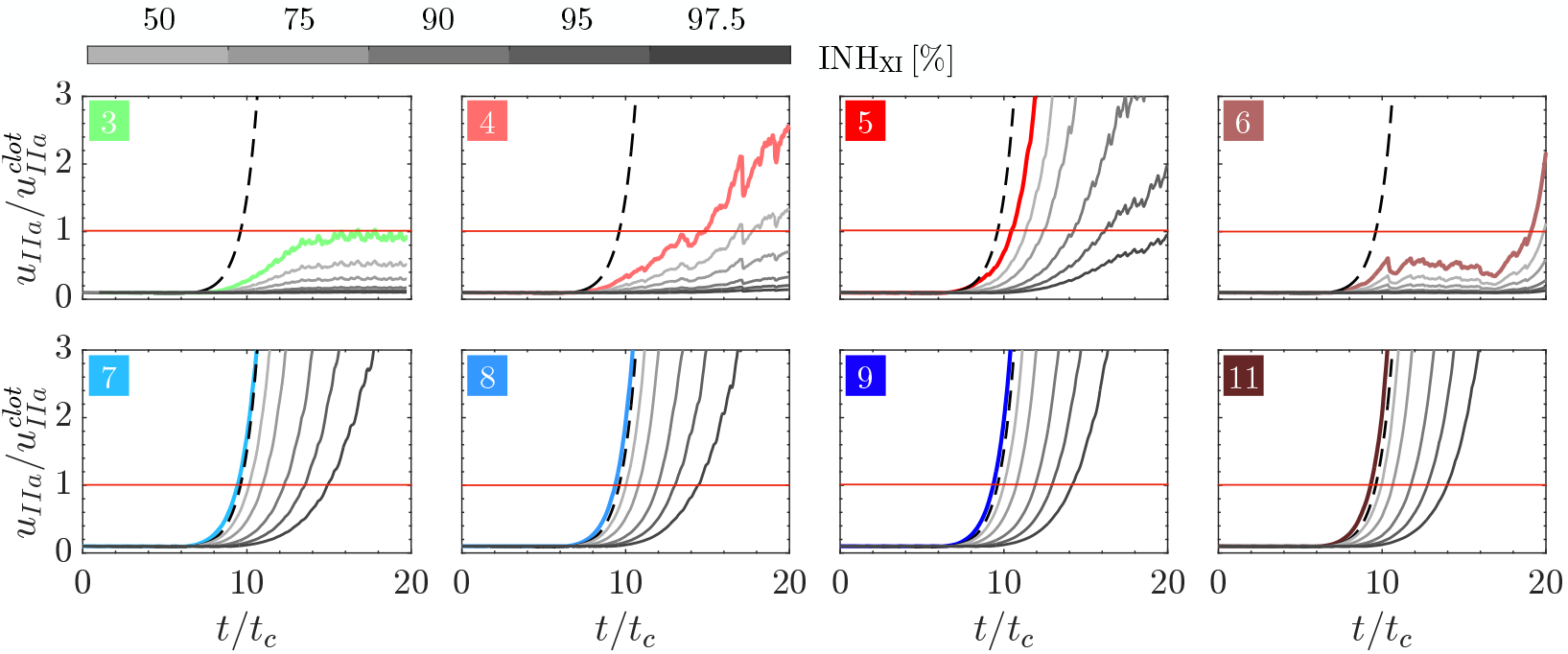
Temporal evolution of maximum thrombin concentration (*u*_*IIa*_) in the LAA across Factor XI Inhibition Level: Time series depicting the maximum thrombin concentration (*u*_*IIa*_) for the nominal case (color lines) and five inhibition levels for factor XI: INH_*XI*_ = [50, 75, 90, 95, 97.5]% 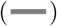 in the LAA of each patient, normalized by the thrombin concentration threshold 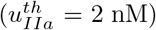. Additionally, the no-flow solution of the 32-ODE system Eq. (9) is provided for reference 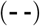.

Figure 10 represents the dynamics of LAA thrombin concentration when inhibiting factor XII, similar to Figure 9. Overall, factor XII inhibition prevented thrombin production more significantly in the *moderately coagulating* LAA than in the *severely coagulating* ones, similar to factor XI. Also, factor XII inhibition was more effective than factor XI inhibition, particularly in the *severely coagulating* group. Of note, Figure 10 suggests that the coagulation threshold was not reached after the 20 cycles of simulation under sufficiently strong factor XII inhibition, even in *severely coagulating* LAAs.

**Figure 10:**
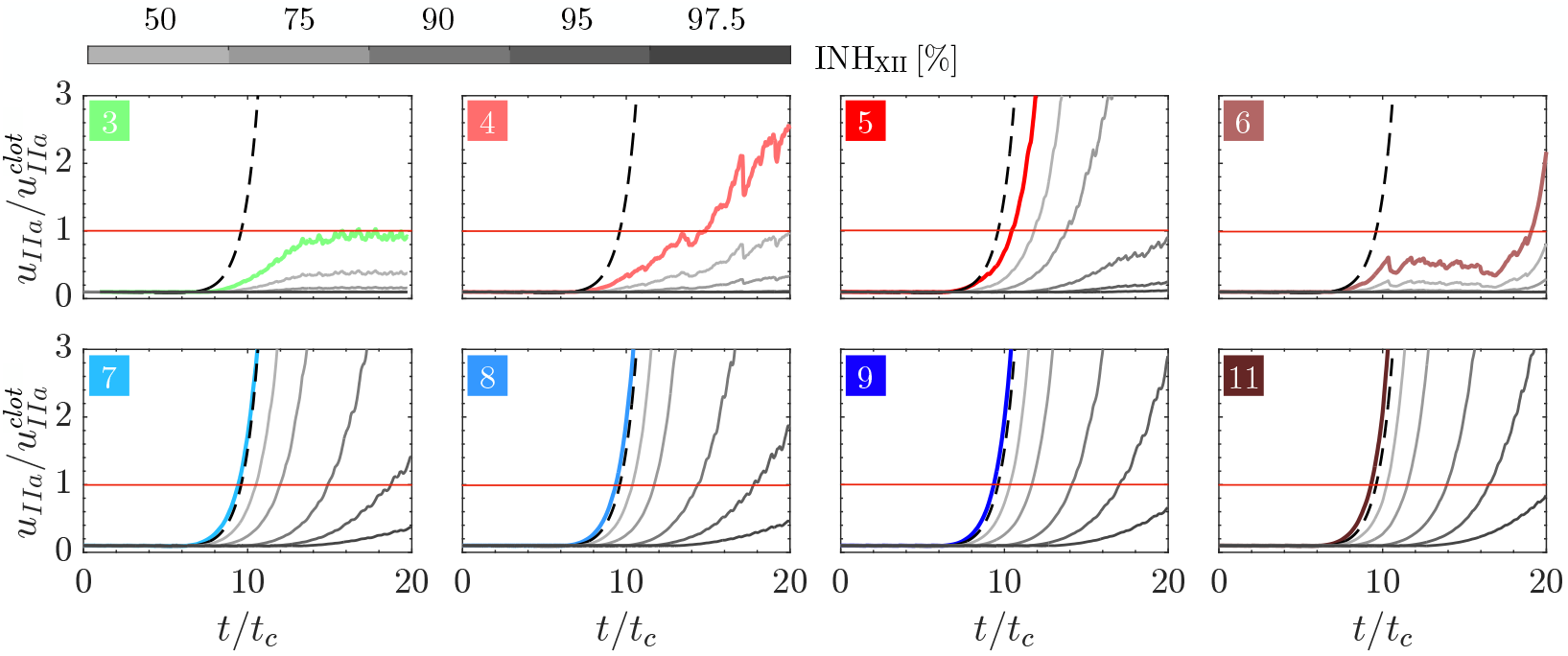
Temporal evolution of maximum thrombin concentration (*u*_*IIa*_) in the LAA across Factor XII Inhibition Level: Time series depicting the maximum thrombin concentration (*u*_*IIa*_) for the nominal case (color lines) and five inhibition levels for factor XII: INH_*XII*_ = [50, 75, 90, 95, 97.5]% 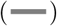 in the LAA of each patient, normalized by the thrombin con-centration threshold 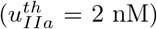. Additionally, the no-flow solution of the 32-ODE system Eq. (9) is provided for reference 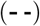.

To quantify more precisely the impact of factor XI and XII inhibition on coagulation, we plotted coagulation time (*t*_*co*_) and coagulating volume (*V*_*co*_) vs. each factor’s INH (Figure 11). In these plots, *V*_*co*_ was averaged over the last simulation cycles and normalized by each patient’s mean LAA volume (*V*_*LAA*_). Figures 11A-B illustrate that after >50% inhibition of factor XI, the *moderately coagulating* cases 4 and 6 did not activate the coagulation cascade within the simulated 20 cycles (i.e., *t*_*co*_ > 20*t*_*c*_ and *V*_*co*_ = 0). In contrast, *severely coagulating* cases 5, 7, 8, 9, and 11 exhibited coagulation times consistent with the no-flow model that were relatively insensitive to factor XI anticoagulation for inhibition levels ≲ 90% (Figure 11A). Consonantly, these cases’ *V*_*co*_ only decreased with anticoagulation for INH_*XI*_ ≳ 90%.

**Figure 11:**
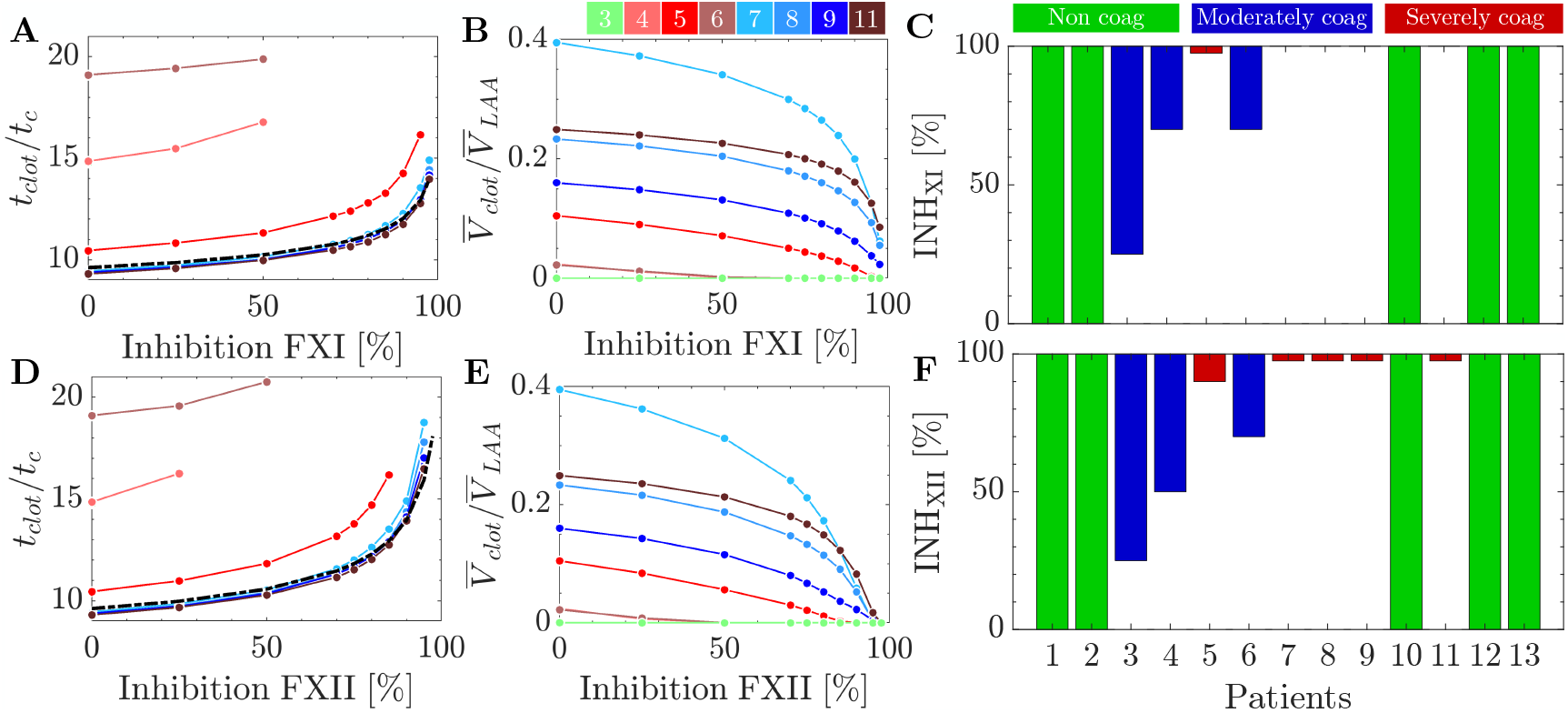
Effect of factor XI and factor XII inhibition levels on coagulation time and volume: (A, B) Coagulation time and mean coagulating volume in the last 5 cardiac cycles (normalized by the mean LAA volume) vs. factor XI inhibition level. (C) Factor XI inhibition range necessary to keep 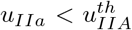. (D, E) Coagulation time and mean coagulating volume in the last 5 cardiac cycles (normalized by the mean LAA volume) vs. factor XII inhibition level. (F) Factor XII inhibition range necessary to keep 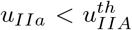. The coagulation time obtained in the no-flow reaction model 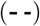 is shown as a reference in panels A and D.

For the most part, inhibiting factor XII had similar effects on *t*_*co*_ and *V*_*co*_ as inhibiting factor XI. However, factor XII anticoagulation was more effective as seen in Figure 11D-E. In particular, the *severely coagulating* cases experienced more dramatic drops in coagulating volume for moderate values of INH_*XII*_ ≈ 75%.

The response to anticoagulation varied among patients as reflected by some of the *V*_*co*_ vs. INH curves crossing each other. For instance, the “untreated” patient with largest *V*_*co*_ in the cohort (case 7) did not have the largest *V*_*co*_ under maximum inhibition of factor XI or factor XII (Figure 11B,E). Figures 11C,F display the range s factor XI/XII inhibition that bring *V*_*co*_ down to zero in each patient. These plots confirm that *non-coagulating* cases did not need inhibition, and show that coagulation can be prevented with moderate levels of factor XI/XII inhibition in *moderately coagulating* cases. They also show that a very significant if not total inhibition of factor XII and especially factor XI is required to stop coagulation in the LAA of *severely coagulating* cases.

Next, we investigated whether each patient’s differential response to anticoagulation was related to LAA blood stasis. Figure 12 depicts the spatial distribution of residence time and thrombin inside a *moderately coagulating* LAA (panel B) and two *severe coagulating* LAAs (panels C and D) after 97.5% inhibition of factors XI/XII. In regions where 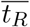 10 *t*_*c*_, both treatments successfully deactivated the coagulation cascade. Blood pools with quasi-perpetual stasis, reaching 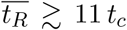 over 20 cycles of simulation, did not respond to 97.5% factor XI inhibition but did respond to a similar inhibition of factor XII. These regions of nearly quasi-perpetual stasis were found in all the *severely coagulating* cases. Particularly, case 11 had areas with 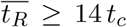 after 20 simulation cycles and had LAA thrombin concentrations very close to the coagulating threshold even after a 97.5% inhibition of factor XII (Figure 12D).

**Figure 12:**
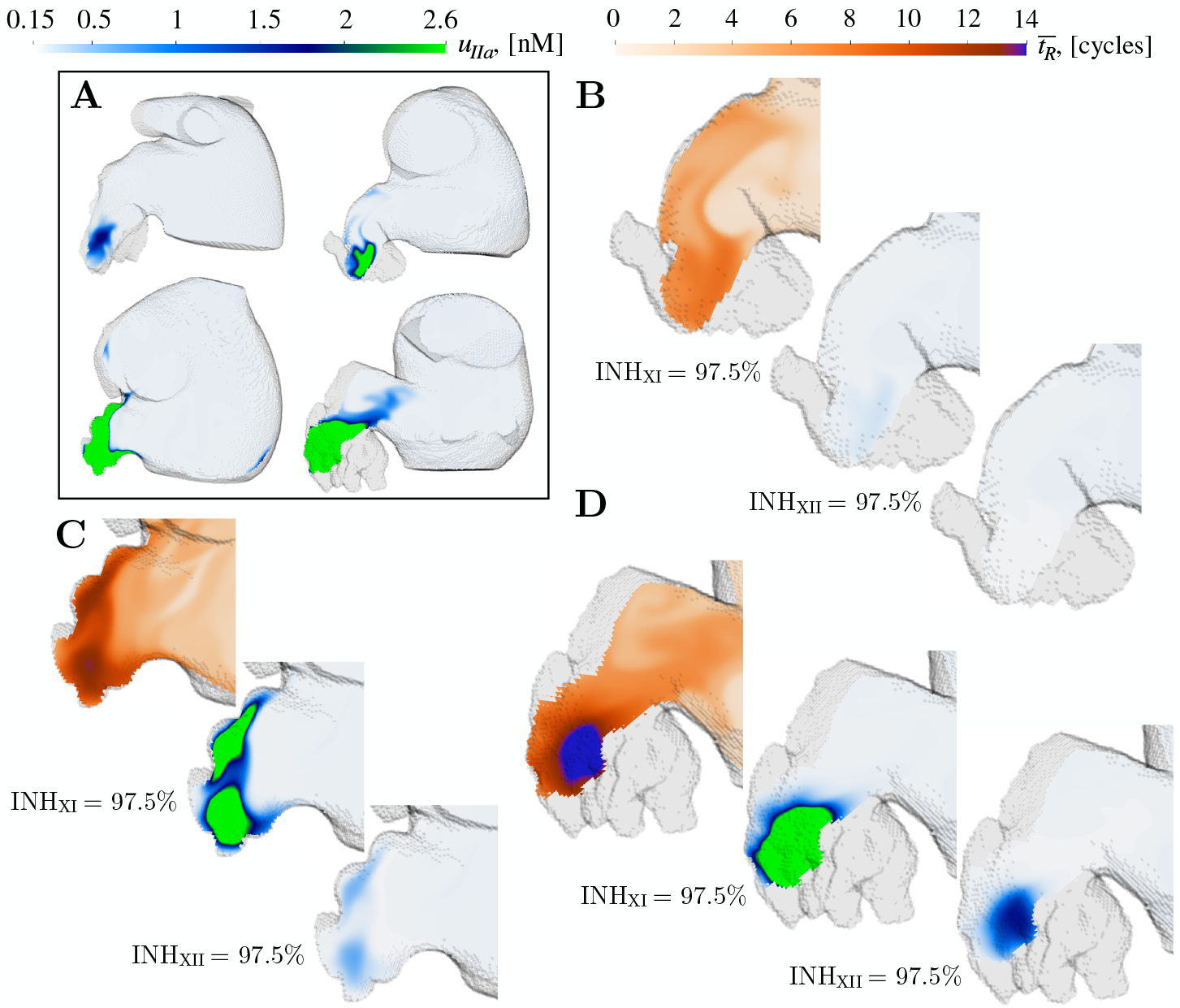
Spatial distribution of residence time 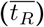 and thrombin accumulation (*u*_*IIa*_) on oblique plane sections after factor XI and XII inhibition: For reference, (A) displays *u*_*IIa*_ for nominal initiation in cases 3,4,7 and 11. Spatial visualization of residence time 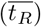 and thrombin accumulation (*u*_*IIa*_) for INH_*XI*_ = INH_*XII*_ = 97.5% on oblique plane sections for Case 4 (B), Case 7 (C) and Case 11 (D) after 20 cardiac cycles *t/t*_*c*_ = 20.

## 4 Discussion

The relevance of left heart flow patterns on thrombosis and cardiogenic stroke has long been recognized [García-Fernández et al., 1992, Delemarre et al., 1990, Martinez-Legazpi et al., 2018]. In recent years, clinical studies are solidifying the causal association between intracardiac stasis and brain embolism [Lee et al., 2014, Costello et al., 2020, Rodríguez-González et al., 2024]. Modeling cardiac thrombosis *in vivo* is challenging because the clotting times measured in humans differ significantly from those measured in commonly used large animal models (e.g., calves, sheep, goats, and pigs) [Mizuno et al., 2018]. Computational models are alternative to experiments but their high computational cost has limited their use, despite noteworthy pioneering efforts in idealized models [Seo et al., 2016, Qureshi et al., 2021]. To the best of our knowledge, there are no systematic simulation campaigns of intracardiac coagulation considering multiple patients, let alone each patient’s response to different anticoagulant doses. To address this paucity, we carried out a wide-ranging series of simulations of left atrial (LA) blood flow and coagulation in 3D patient-specific anatomies, testing different levels of factor XI/XII inhibition.

### Multi-Fidelity models accelerate patient-specific simulations of left atrial coagulation by two orders of magnitude

Mathematical models of the coagulation cascade are often formulated as systems of ordinary differential equations (ODEs) representing the cascade’s reaction kinetics [Cito et al., 2013] These ODE systems are valid when the coagulation species form a homogeneous mixture throughout the volume of interest, but intracardiac flow patterns create regions with distinct transport that impede homogeneous mixing [Bermejo et al., 2015, Eriksson et al., 2013]. Therefore, advection-reaction-diffusion partial differential equations are required to model coagulation inside the cardiac chambers, and solving these PDEs in three-dimensions is computationally intensive, especially as the number of coagulation species increases.

Multi-fidelity (MuFi) coagulation modeling is rooted in the observation that the reaction terms in the PDEs governing species concentration can be evaluated independently at each spatial point as long as there is no mass diffusivity, so that these equations can be converted into ODEs [Fogelson and Guy, 2008]. This idea has been recently formalized and extended to non-zero diffusivities by Taylor-expanding the ODEs around the zero-diffusivity limit, producing spatiotemporal maps of species concentrations in terms of the statistical moments of residence time [Guerrero-Hurtado et al., 2023]. The MuFi approach reduces the problem of solving *N* reaction-advection-diffusion for *N* coagulation species to *p* PDEs for the first *p* statistical moments of 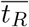 and *N* ODEs for the reaction kinetics. Therefore, if a MuFi-*p* model reproduces the high-fidelity (HiFi) reaction-advection-diffusion results for a given cardiovascular geometry, reaction kinetics, and sufficiently low order *p*, MuFi modeling accelerates the simulation of blood coagulation under flow for that configuration.

In this work, the effectiveness of the MuFi models was verified in patient-specific LA flows by comparing it to a 9-species HiFi PDE coagulation cascade model with reaction kinetics described in Zarnitsina et al. [2001]. Due to the elevated computational cost of the HiFi simulations, we restricted the verification to two distinct patient-specific LA flows: one corresponding to a subject with normal LA function and sinus rhythm and another with impaired LA function and atrial fibrillation (AF). In both cases, a second-order MuFi model (MuFi-2) was sufficient to capture the spatiotemporal dynamics of thrombin over 20 cardiac cycles with errors in thrombin concentration lower than 10%.

There are two ways by which MuFi models speed up simulations of the coagulation cascade in flowing blood. First, running one instance of a given coagulation model is predicted to be *αN/p* times faster in MuFi-*p* form than in HiFi form, where *α* ≳ 1 is a proportionality constant [Guerrero-Hurtado et al., 2023]. Second, and more important, MuFi models decouple the CFD solver from the coagulation solver, providing countless virtually free coagulation cascade simulations per patient-specific CFD simulation. Therefore, running a series of *k* coagulation cascade simulations on one patient can achieve a speedup of ∼ *kN/p*, which grows boundlessly with the size of the campaign and the number of coagulation species.

As an example, let us consider the 32-species model used for the MuFi-2 simulation campaign in this study. Extrapolating the speedup values of 7.7 and 4.4 obtained running MuFi-2 and MuFi-3 for the 9-species model [Zarnitsina et al., 2001], *αN/p* ≈ 25 is a reasonable estimate for the 32-species model. Then, since each patient-specific inhibition study reported in this manuscript involved *k* = 19 MuFi-2 coagulation simulations (no inhibition plus 9 inhibitions of Factors XI and XII), the cumulative speedup of the whole simulation campaign would be ≈ 19 *×* 25 = 475. It is difficult to conceive the simulations reported in this manuscript as feasible should they require 475X compute time.

### Atrial hemodynamics, coagulation cascade, and anticoagulation therapy

AF causes weak, irregular atrial beats, producing abnormal flow patterns that can lead to thrombosis and increasing the risk of stroke by a factor of 5 [Wolf et al., 1991]. Patients diagnosed with AF often need long-term anticoagulation treatment to lower the risk of ischemic stroke and related embolic events. Selecting the appropriate treatment and dosage for an individual patient is a complex decision that involves balancing the treatment benefit with its bleeding risk [Joglar et al., 2024]. Anticoagulant agents targeting the intrinsic cascade, such as factor XI inhibitors, may significantly decrease bleeding risk as indicated by recently completed phase-II randomized clinical trials (RCT)[Piccini et al., 2022]. However, the interruption of some phase-III RCTs for these agents suggests an incomplete understanding of their efficacy vs. dosage [Presume et al., 2024]. Because atrial clotting is sensitive to patient-specific flow patterns, these patterns are also likely to affect each patient’s response to anticoagulation therapy. This dependence, and the increasing availability of anticoagulation drugs with different mechanisms of action, warrant investigating the interplay between AF, atrial hemodynamics, and coagulation in patient-specific models.

We simulated the activation of the coagulation cascade in atrial blood under different levels of factor XI and factor XII inhibition in *N* = 13 patient-specific LA models. In all patients studied, no-flow models based on ODEs severely overpredicted thrombin concentration, confirming that flow patterns play a significant role in LA thrombosis. The 3D snapshots of thrombin concentration showed that it peaks within the LAA, consistent with the LAA being the most common thrombotic site and having highest blood residence time 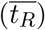 across the LA [Zhang and Gay, 2008, Koizumi et al., 2015, Markl et al., 2016, García-Isla et al., 2018, D’Alessandro et al., 2020, Sanatkhani et al., 2021, Zhang et al., 2023, Zingaro et al., 2024, Bäck et al., 2023, Bshennaty et al., 2024, Musotto et al., 2024]. The spatiotemporally resolved distributions of coagulation species concentrations were analyzed to derive global coagulation metrics, such as the maximum thrombin concentration, coagulation time (*t*_*co*_), and coagulating volume (*V*_*co*_). The maximum thrombin concentrations indicated different levels of coagulation cascade activation in 8 out of the 13 patients, whereas no activation was detected in the remaining 5 patients. We defined the former as *coagulating* cases and the latter as *non-coagulating* cases. Among the *coagulating* cases, we observed two distinct patterns in the temporal dynamics of maximum thrombin concentration and coagulating volume. The *severely coagulating* cases showed exponential growth of the thrombin concentrations, while the *moderately coagulating* cases showed more complex dynamics with significantly slower net growth and smaller coagulating volumes.

The characteristics of our patient cohort, discussed in the limitations section below, prevent a rigorous statistical analysis of our models’ predictive value. Yet, it was encouraging to see that all cases identified as *severely coagulating* in our model had an AF diagnosis, were imaged and therefore simulated while in AF, and had severely impaired atrial function (emptying fraction *EF* ≲ 0.13). Consonantly, large portions of these patients’ LAAs had recirculating regions that did not exchange fluid with the rest of the atrium. All *noncoagulating* cases except one were negative for AF and LAA thrombus and had relatively normal atrial function. The exception was an AF patient imaged and simulated in sinus rhythm, with the highest EF among the AF group (0.26). The *moderately coagulating* cases had mixed risk factors. One had a normal atrial function but a relatively large LAA. The others were imaged in AF and had moderately impaired atrial function (*EF* ≈ 0.23). Of note, all the patients with LAA clot or a history of TIAs had *moderately* or *severely coagulating* hemodynamic substrates.

Inhibiting either factor XI or factor XII blunted thrombin growth in our models, although factor XII seemed more effective than factor XI for the same inhibition level. *In vitro* and *in vivo* experiments [Wilbs et al., 2020, Liu et al., 2011, Naito et al., 2021] indicate that inhibition levels between 70% and 90% of the target factor can prolong aPTT and lower thrombin concentrations significantly. Our results are generally consistent with these experiments and suggest that the efficacy of target inhibition is patient-specific. A modest inhibition of either factor (INH_*i*_ < 50%) was sufficient to arrest thrombin growth in the *moderately coagulating* cases. In contrast, the *severely coagulating* cases required drastic factor XII inhibition (INH_*XII*_ = 97.5%) to arrest thrombin growth. Furthermore, 97.5% inhibition of factor XI was unable to bring *V*_*co*_ down to zero.

Since we used the same reaction kinetics for all patients, the differences in coagulation dynamics and anticoagulant efficacy can be attributed to differences in patient-specific LA hemodynamics. In particular, to the spatiotemporal distributions of the residence time inside the LAA, which quantifies LAA blood stasis [Sanatkhani et al., 2021, Rigatelli et al., 2023, Bäck et al., 2023, Fang et al., 2022, Paliwal et al., 2024]. The view emerging from recent simulation studies is that LAA blood stasis depends on multiple factors including LA function [Garcia-Villalba et al., 2021, Gonzalo et al., 2022, 2024], PV position, orientation and/or flow split [García-Isla et al., 2018, Dueñas-Pamplona et al., 2022, Durán et al., 2023, Musotto et al., 2024], and LAA morphology [Bosi et al., 2018, Masci et al., 2019, Grigoriadis et al., 2020, Morales Ferez et al., 2021]. In addition to confirming that 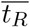 is associated with LAA thrombosis risk, our simulations indicate that the efficacy of anticoagulation therapy may also depend on LAA 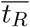. Considering this evidence, the optimal anticoagulant agent and dosage could vary significantly from patient to patient. Moreover, our results suggest that anticoagulation alone might be insufficient to prevent clotting in regions with quasi-perpetual stasis. This result is consistent with the low albeit positive rates of ischemic stroke observed in anticoagulated AF patients [Seiffge et al., 2020], and suggests that LAA occlusion might be preferrable to anticoagulation in patients with extremely slow flow in the LAA. The computationally efficient tools presented in this manuscript lay the foundation to personalize the assessment of coagulation risk and anticoagulation therapy based on medical imaging. These tools could also be applied to refine patient selection for clinical trials of novel anticoagulation drugs.

### Study limitations

This study’s patient group, *N* = 13, is significantly larger than the *N* = 2 used in the only other LA clotting simulation study we know of [Qureshi et al., 2023]. However, it is still far from sufficient to achieve statistical power. Moreover, our patient selection prioritized achieving a wide range of atrial functions and volumes and over-represented AF and LAA thrombosis to demonstrate the interplays between AF, LA hemodynamics, and thrombosis. For these reasons, while we found interesting trends of agreement between our model results, e.g., the *non-coagulating, moderately coagulating*, or *severely coagulating* class, and patient characteristics, e.g., presence of LAA clot or TIAs, our data is insufficient to ascertain whether these trends indicate correlation.

We note but are less concerned about the subjectivity of classifying patients as *non-coagulating, moderately coagulating*, or *severely coagulating*. Given that the definition of coagulation time is not unique and the thrombin threshold values used to identify clotting vary in the range of 2-15 nM [Greenberg et al., 1985, Danforth et al., 2012, Kogan et al., 2001], our classification may seem somewhat subjective (exponential growth vs. significantly slower growth; maximum thrombin concentration over threshold value 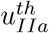), but our data suggest it is reasonably robust. If we lowered or raised this threshold concentration by a factor of 2, only one case would switch class between *non-coagulating* to *moderately coagulating*. And changing the definition of sufficiently fast growth for the *severely coagulating* group might switch only one case from *moderately* to *severely coagulating*, if any.

Although our CFD simulations used patient-specific LA shapes and patient-specific LA motion obtained from 4D-CT imaging, several parameters in our models were not patient-specific. Below, we discuss these parameters and the potential limitations of using generic values across the patient population. In all cases, the principal reason for using generic values was the lack of data either in direct form or in a form that would allow for identifying parameters in our models. Considering this lack and that our main objective was to evaluate the effect of factor XI/XII inhibition on LA thrombosis, it seemed sound to systematically vary the initial concentrations of factor XI/XII while keeping all other parameters constant.

All simulations were run at a constant heart rate (60 bpm), and the PV flow rates were evenly split to set inflow/outflow boundary conditions based on each patient’s LV and LA time-dependent volumes, as in Garcia-Villalba et al. [2021]. While prior studies justify this choice [Lantz et al., 2019], the PV flow split could affect flow patterns inside the LAA, par-ticularly in cases where LAA residence time is high and thrombosis is more likely [Durán et al., 2023]. We considered non-Newtonian blood rheology, but we fixed the hematocrit value (*Hct* = 43.5) because this parameter was unavailable for some patients. We also fixed the characteristic time of RBC aggregation (*t*_*µ*_ = 3 s) as described in Gonzalo et al. [2022]. These two parameters could impact residence time and non-Newtonian effects in the LA [Vasudevan et al., 2017]. The reaction kinetics in the coagulation cascade model were not patient-specific either, using the same reaction rates and initial conditions for all cases. Variations in the central blood concentration of coagulation factors and equilibrium constants can be significant [Ratto et al., 2021]. We did not have patient-specific measurements of these concentrations or clotting times, and even if these data were available, many parameters of our detailed 32-species kinetic model would not be identifiable in a patient-specific manner.

We only considered the intrinsic coagulation pathway since our main focus was factor XI/XII inhibition, ignoring contributions from the extrinsic pathway that could influence thrombin generation [Davie et al., 1991]. Due to the high computational cost of running HiFi simulations in 3D patient-specific anatomies, we verified our MuFi approach vs. the HiFi approach using a 9-species model [Zarnitsina et al., 2001], then ran our simulation campaign using a 32-species model [Zhu, 2007]. Finally, we adopted an oversimplistic representation of anticoagulation threapy by directly varying the inhibition level of each target factor (INH_*i*_). In reality, clinicians can regulate anticoagulant dose, but controlling INH_*i*_ is more challenging due to numerous factors such as drug absorption, distribution, metabolism, and elimination [Hindley et al., 2023].

## 5 Conclusions

We applied multi-fidelity (MuFi) coagulation cascade modeling to 3D patient-specific left atrial segmentations with hemodynamics obtained from computational fluid dynamics. We considered a detailed model of the intrinsic coagulation pathway sweeping over a wide range of factor XI/XII inhibition levels in patients with varying LA anatomy and function, sinus and atrial fibrillation, and with and without LAA thrombosis. We ran a total 247 simulations, which to the best of our knowledge, constitutes the first systematic study of intracardiac coagulation and anticoagulation therapy using 3D patient-specific anatomies and hemodynamics.

We found that thrombin exhibited the most significant growth in the LAA of patients with impaired blood washout. Our results also suggest that the effectiveness of novel an-ticoagulation agents targeting the intrinsic coagulation pathway in AF may depend on patient-specific flow patterns. By providing computationally efficient tools to study this dependence, this work lays the foundation for improving patient selection for clinical trials and personalizing the prescription of anticoagulant agents based on medical imaging.

## Appendix

### A1. Coagulation cascade for the verification runs

In this section, we describe the reaction terms for the 9-species coagulation model employed in the valudation study. The model, initially introduced by Zarnitsina et al. [2001], incorporates three positive feedback loops, which are facilitated by thrombin-activated factors XIa, Va, and VIIIa. Additionally, it includes negative feedback mechanisms, where factors Va and VIIIa are deactivated by the generation of PCa, which is itself activated by thrombin. The model was validated against experiments by Ataullakhanov et al. [1994] for various calcium concentrations. The reaction rates for the nine species are detailed in eqs. (18-27), and the reaction coefficients reported by Ataullakhanov et al. [1994] are listed in Table 3.

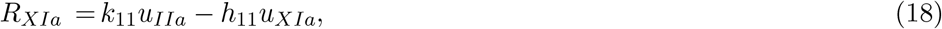

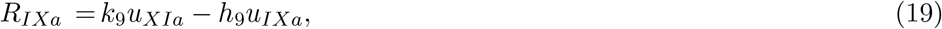

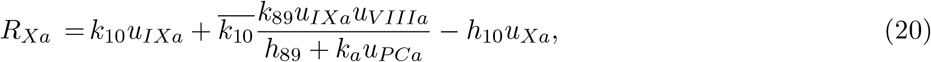

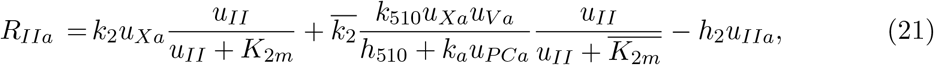

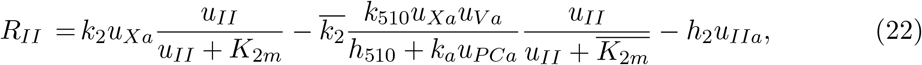

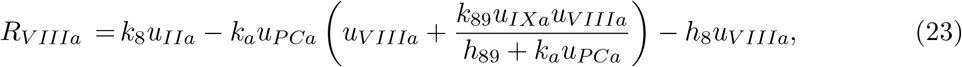

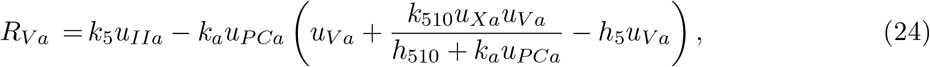

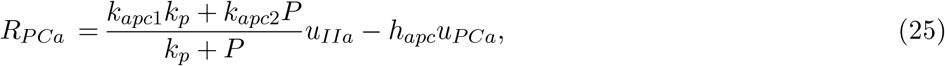

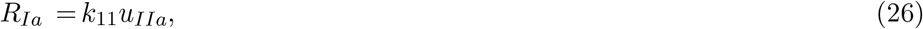

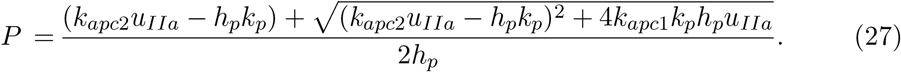

**Table 3:**
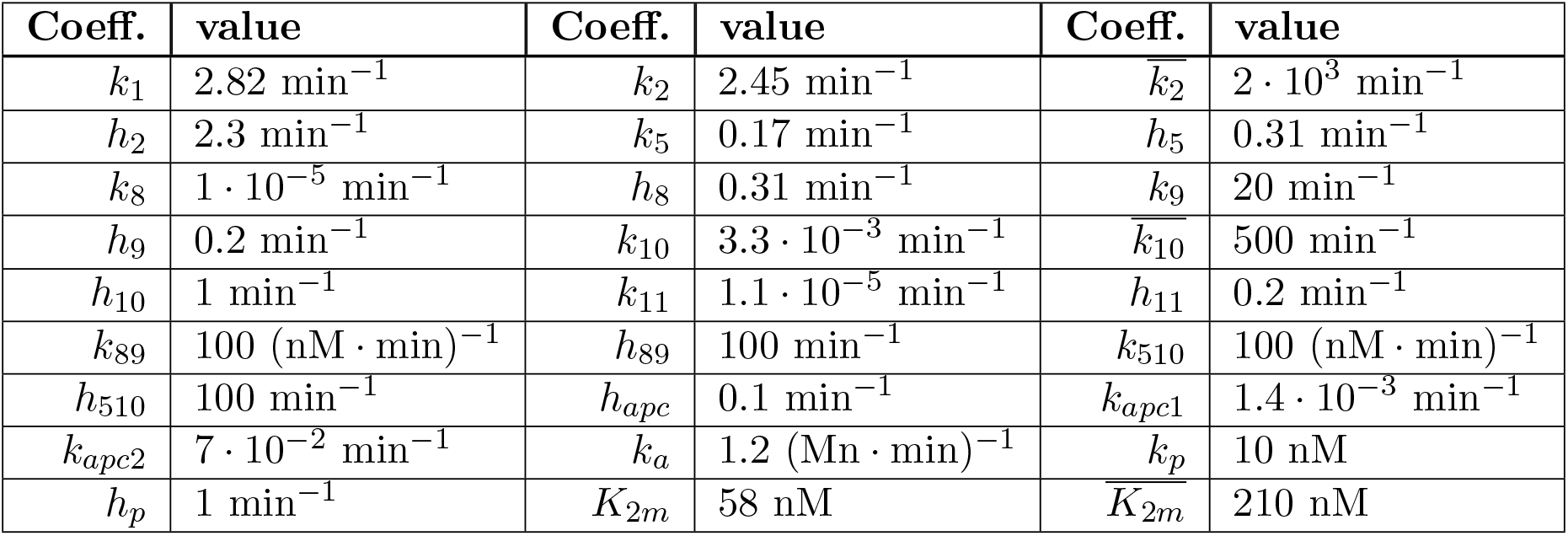
Reaction Rates.

